# Brain-wide Correspondence Between Neuronal Epigenomics and Long-Distance Projections

**DOI:** 10.1101/2023.05.01.538832

**Authors:** Jingtian Zhou, Zhuzhu Zhang, May Wu, Hanqing Liu, Yan Pang, Anna Bartlett, Angeline Rivkin, Will N. Lagos, Elora Williams, Cheng-Ta Lee, Paula Assakura Miyazaki, Andrew Aldridge, Qiurui Zeng, J.L. Angelo Salinda, Naomi Claffey, Michelle Liem, Conor Fitzpatrick, Lara Boggeman, Zizhen Yao, Kimberly A. Smith, Bosiljka Tasic, Jordan Altshul, Mia A. Kenworthy, Cynthia Valadon, Joseph R. Nery, Rosa G. Castanon, Neelakshi S. Patne, Minh Vu, Mohammad Rashid, Matthew Jacobs, Tony Ito, Julia Osteen, Nora Emerson, Jasper Lee, Silvia Cho, Jon Rink, Hsiang-Hsuan Huang, António Pinto-Duartec, Bertha Dominguez, Jared B. Smith, Carolyn O’Connor, Hongkui Zeng, Kuo-Fen Lee, Eran A. Mukamel, Xin Jin, M. Margarita Behrens, Joseph R. Ecker, Edward M. Callaway

**Author notes:** These authors contributed equally (Jingtian Zhou, Zhuzhu Zhang).

## Abstract

Single-cell genetic and epigenetic analyses parse the brain’s billions of neurons into thousands of “cell-type” clusters, each residing in different brain structures. Many of these cell types mediate their unique functions by virtue of targeted long-distance axonal projections to allow interactions between specific cell types. Here we have used Epi-Retro-Seq to link single cell epigenomes and associated cell types to their long-distance projections for 33,034 neurons dissected from 32 different source regions projecting to 24 different targets (225 source →target combinations) across the whole mouse brain. We highlight uses of this large data set for interrogating both overarching principles relating projection cell types to their transcriptomic and epigenomic properties and for addressing and developing specific hypotheses about cell types and connections as they relate to genetics. We provide an overall synthesis of the data set with 926 statistical comparisons of the discriminability of neurons projecting to each target for every dissected source region. We integrate this dataset into the larger, annotated BICCN cell type atlas composed of millions of neurons to link projection cell types to consensus clusters. Integration with spatial transcriptomic data further assigns projection-enriched clusters to much smaller source regions than afforded by the original dissections. We exemplify these capabilities by presenting in-depth analyses of neurons with identified projections from the hypothalamus, thalamus, hindbrain, amygdala, and midbrain to provide new insights into the properties of those cell types, including differentially expressed genes, their associated cis-regulatory elements and transcription factor binding motifs, and neurotransmitter usage.

## Introduction

In any given brain, each neuron contributes uniquely to brain function. Nevertheless, neurons can be grouped into types based on similarities and differences across multiple dimensions, including epigenetic state, gene expression, anatomy, and physiology. Single-cell genomic technologies have been particularly impactful for cell type classification due to their high throughput (millions of cells assayed) and dimensionality (thousands of genes and even more genetic loci) leading to the identification of large numbers of transcriptomic and epigenomic clusters corresponding to possible cell types across the entire mouse brain.

A prominent and distinguishing anatomical feature of many brain neuron types is their long-distance axonal projections. Long-distance projections can be directly related to single neuron gene expression or epigenomes by use of powerful linking technologies, including BARseq^1,2^, Retro-seq^3,4^, and Epi-Retro-Seq^5^. Previous studies have used Retro-Seq and Epi-Retro-Seq to link mouse neocortical^3–5^, hypothalamic^6^ and thalamic projection cell types^7^ to their genetic and epigenetic clusters, revealing complex but predictable relationships. For example, cortical neurons projecting solely to intra-telencephalic (IT) targets fall into different clusters than those that project to extra-telencephalic (ET) targets. On the other hand, cortical layer 2/3 (L2/3) IT neuron types projecting to different cortical areas typically co-cluster despite having quantifiable and predictable genetic and epigenetic differences across the populations^4,5^. In the face of this complexity, how can single-cell genetic and epigenetic assays be used to inform the structure and function of brain cell types and how can neuronal structure predict genetics, epigenetics, and function? Further, can the principles learned from more limited previous studies be extended to the entire brain? Are there different principles linking projection status to epigenetics for different brain areas?

To address these questions we employed Epi-Retro-Seq to assay 33,034 neurons from 225 source →target combinations across the entire mouse brain. This approach combines retrograde labeling with single nucleus methylation sequencing (snmC-Seq), which allows identification of potential gene regulatory elements and prediction of gene expression in the same neuron. Gene expression can be predicted because non-CG (CH; H=A,T,C) methylation of gene bodies is inversely related to RNA expression^8,9^, while epigenetic elements regulating expression can be identified using methylation at CG (mCG) dinucleotides^8^. It is also expected that Epi-Retro-Seq can provide unique insight into developmental mechanisms that shape connectivity because CH methylation accumulates during and peaks at the end of the developmental critical period, and CG methylation is reconfigured during synaptic development^10^.

## Results

### Epi-Retro-Seq of 225 brain-wide projections

To link single-neuron epigenomes to their projection targets and cell body locations, we used Epi-Retro-Seq^5^. A retrogradely-infecting AAV vector expressing cre-recombinase (AAV-retro-Cre^11^) was injected into the brains of cre-dependent, nuclear-GFP expressing reporter mice (INTACT-cre^8^) at a target region of interest (**Fig. 1a**). Four mice (2 male and 2 female) were injected for each of 24 different target brain areas, including targets in the isocortex (CTX), hippocampal formation (HPF), olfactory areas (OLF), amygdala (AMY), cerebral nuclei (CNU), interbrain (IB), midbrain (MB), hindbrain (HB), and cerebellum (CB) (**Fig. 1a,b, Supplementary Table 1**). After 2 weeks, mice were sacrificed and the brain was hand dissected following the Allen Mouse Common Coordinate Framework (CCF), Reference Atlas, Version 3^12^, into 32 possible source regions spanning the same major brain structures as the target injections (**Fig. 1a,b and Extended Data Fig. 1**). For any given mouse, dissected sources corresponding to locations with known projections to the target were selected for profiling. Nuclei preps were made from dissected source tissue and subject to fluorescence-activated nuclear sorting (FANS) for GFP-positive, NeuN-positive retrogradely labeled neuronal nuclei which were then processed for single nucleus methylation sequencing (snmC-seq; **Fig. 1a, Methods**)^13–15^.

**Fig 1.**
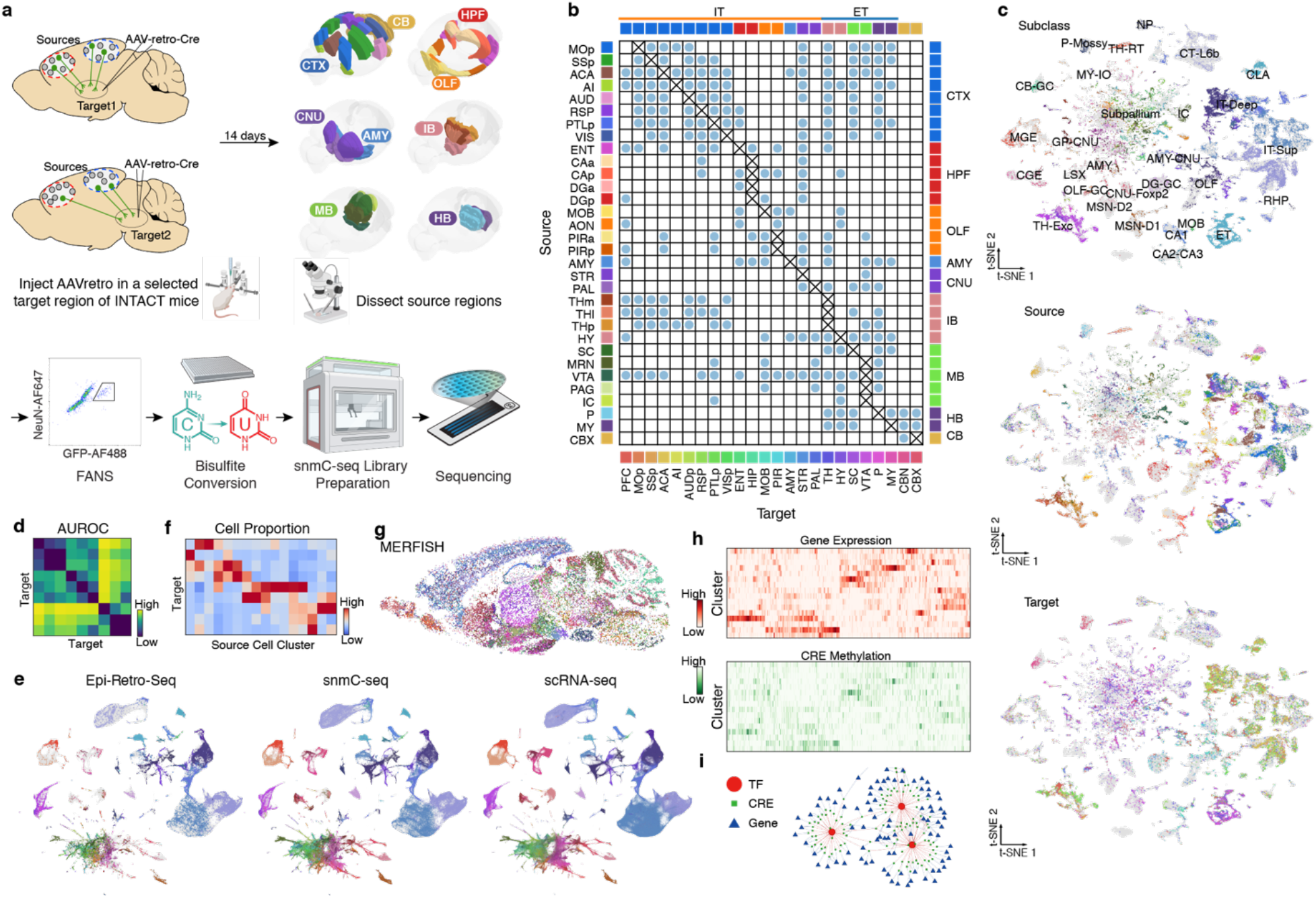
The epigenomic landscape of brain-wide projection neurons. **a**, Schematics of Epi-Retro-Seq workflow for retrogradely labeling and epigenetically profiling single projection neurons. The retrograde tracer rAAV2-retro-cre was injected into a specific target area in INTACT knock-in mice to label the nuclei of neurons that project to the target area with cre-dependent nuclear GFP. Source regions of interest with projections to the target areas were dissected 14 days after injection. Single GFP+/NeuN+ nuclei were isolated using fluorescence activated nuclei sorting (FANS), and then subjected to snmC-seq library preparation and sequencing for epigenome profiling. Brain diagrams of the source regions were derived from the Allen Mouse Brain Reference Atlas (version 3 (2020)). **b**, 225 source target combinations were profiled using Epi-Retro-Seq from 32 different source regions projecting to 24 different targets across the whole mouse brain. **c**, Joint two-dimensional t-distributed stochastic neighbor embedding (t-SNE) of Epi-Retro-Seq (n=35,938) and unbiased snmC-seq (n=276,187) neurons. snmC-seq neurons are in gray and Epi-Retro-Seq neurons are colored by cell subclass (top), the source regions of neurons (middle, same color palette as row colors on the left of (**b**)), or their projection targets (bottom, same color palette as column colors on the bottom of (**b**), n=33,034, after removing the experiments with less confident target assignment). **d**, As an example, area under the curve of receiver operating characteristic (AUROC) for pairwise comparisons of amygdala neurons projecting to 9 targets. Higher AUROC scores suggest greater distinguishability between the compared projections based on their gene body CH methylation (mCH) levels. **e**, Joint t-SNE of whole mouse brain neurons from Epi-Retro-Seq (n=35,743), unbiased snmC-seq (n=266,740), and single-cell RNA-seq (scRNA-seq, n=2,434,472) colored by cell subclass. **f**, As an illustration, the proportion of neurons found in each amygdala cell cluster (row) that projects to each target (column). Only clusters that were enriched for projection neurons are shown and values are Z-score normalized across targets. **g**, A sagittal brain slice for multiplexed error robust fluorescence in-situ hybridization (MERFISH) with all neurons colored by their assigned subclasses. **h**, An illustration of joint analysis of single-cell transcriptomes and DNA methylomes that enables the characterization of gene expression patterns of differentially expressed genes (DEGs) between these projection-enriched clusters, as well as the CG methylation (mCG) levels of DEG-associated putative cis-regulatory elements (CREs), as marked by differentially methylated regions (DMRs). **i**, An illustration of identifying transcription factors (TFs) whose binding motifs are enriched in these CREs and potentially regulate the expression of the DEGs.

After basic quality control, we recovered 48,032 single-cell methylomes which were mapped to an unbiased sample of snmC-seq data with 301,626 cells (companion paper #6) to perform cell type classification, and for removal of potential doublets **(Extended Data Fig. 2a-e, Methods**). Each single neuron in the Epi-Retro-Seq sample was assigned to one of the 2,304 level 4 clusters (L4 type) identified in our companion study (companion paper #6). We have previously described cortical neurons from the same 8 cortical sources included here and projecting to 4 cortical and 6 subcortical targets (63 combinations)^5^. For cortical sources, we now incorporate data for an additional 5 cortical targets and 2 more subcortical targets. Similar to our previous work, for cells from cortical sources we included an additional quality control step to eliminate experiments with inadvertent spread of injected AAVretro into source regions (e.g. from AAV spread along injection pipette paths extending through some cortical regions) or with poor quality FANS sorting, by filtering based on the proportion of known on-target vs off-target cell types. (Such filtering is not needed for deeper source regions that are not traversed by injection pipettes.) In total, 33,034 single-nucleus methylomes were analyzed from 225 source →target combinations for which the projection target could be confidently assigned. These neurons were mapped to the unbiased snmC-seq dataset to visualize the epigenetic similarity of projection neurons across cell subclasses, sources, and targets (**Fig. 1c**).

#### Data analysis approaches and visualization across the whole brain

Overarching questions that can be addressed by this large data set include: how distinct are neurons from a given source that project to different targets? And are neurons in different sources that project to the same target combinations more or less distinguishable? To provide a resource that can be used to address the distinguishability of neurons with different projection targets, we quantified which projection types are epigenetically more different than the others by computing area under the curve of receiver operating characteristic (AUROC) for each of the target pairs from every source region (926 pairwise comparisons in total; **Fig. 1d and Extended Data Fig. 3**). An example of the results from the pairwise comparisons for amygdala neurons projecting to 9 targets is shown in **Figure 1d**. Such plots allow visualization of similarities and differences between source neurons projecting to different targets, as further exemplified in our deeper analyses of selected source regions within the main text and figures below. Similar insights can be gained for projection neurons from all of the assayed sources by accessing the complete set of 926 AUROC comparisons (**Extended Data Fig. 3d**).

To facilitate further, comprehensive multimodal characterization of projection neuron types, we integrated the Epi-Retro-Seq data with unbiased samples of snmC-seq described above, and single-cell RNA-seq (scRNA-seq) data containing 2.6 million neurons from 87 micro-dissected brain regions (**Fig. 1e and Extended Data Fig. 4**). Alignment of Epi-Retro-Seq data to these larger and carefully annotated datasets allows for the confident assignment of our cells to consensus clusters and enables the use of consistent nomenclature to describe the correspondence between projection targets and cell types/clusters. We performed co-clustering of the three datasets to identify the cell clusters associated with each projection type, which allows for identification of projection-enriched clusters (see further below). Quantification of the proportion of cells found in each enriched cluster that projects to each target is illustrated for amygdala source neurons in **Figure 1f**. This approach exemplifies analyses explored in detail for other sources, below, and which are provided for all of the sources in the data set in **Extended Data Figure 5**.

To separate neurons projecting to particular targets from different sources, we performed microdissections of freshly cut brain slices. While these careful dissections effectively separate fairly small structures, most dissected regions contain still smaller known anatomical regions, as typically illustrated in mouse brain atlases (**Extended Data Fig. 1**). To potentially link projection-enriched clusters from particular sources to more precise anatomical loci, we performed further integration with multiplexed error robust fluorescence in-situ hybridization (MERFISH) data, allowing examination of the spatial locations of the cells belonging to particular clusters (**Fig. 1g and Extended Data Fig. 6**). Joint atlasing of single neuron transcriptomes and epigenomes further allowed analyses of both the signature genes in projection-enriched clusters based on RNA expression, and methylation profiles to identify differentially methylated regions (DMRs) as putative cis-regulatory elements (CREs) and transcription factors whose binding motifs are enriched in these DMRs (**Fig. 1h,i**).

We prepared extended data figures to allow visualization of the integrative analysis approaches described above (e.g. **Fig. 1d-i**) for all source →target combinations in our dataset (**Extended Data Figs. 3-6**). These integrative analyses were facilitated by combining source regions from the whole brain datasets into 12 larger “region groups” that were common to all 3 data modalities, prior to integration (**Extended Data Fig. 2f,g**). The groups include isocortex (CTX), retro hippocampal region (RHP), piriform area (PIR), hippocampal region (HIP), main olfactory bulb and anterior olfactory nucleus (MOB+AON), striatum (STR), pallidum (PAL), amygdala (AMY), thalamus (TH), hypothalamus (HY), midbrain (MB), and hindbrain (HB).

CTX, isocortex; CB, cerebellum; OLF, olfactory areas; HIP, hippocampal region; CNU, cerebral nuclei; AMY, amygdala; IB, interbrain; MB, midbrain; HB, hindbrain; PFC, prefrontal cortex; MOp, primary motor cortex; SSp, primary somatosensory cortex; ACA, anterior cingulate cortex; AI, agranular insular cortex; AUD, auditory cortex; AUDp, primary auditory cortex; RSP, retrosplenial cortex; PTLp, posterior parietal cortex; VIS, visual cortex; VISp, primary visual cortex; ENT, entorhinal cortex; CAa, anterior Cornu Ammonis; CAp, posterior Cornu Ammonis; DGa, anterior dentate gyrus; DGp, posterior dentate gyrus; MOB, main olfactory bulb; AON, anterior olfactory nucleus; PIR, piriform cortex; PIRa, anterior piriform cortex; PIRp, posterior piriform cortex; STR, striatum; PAL, pallidum; TH, thalamus; THm, anterior medial thalamus; THl, anterior lateral thalamus; THp, posterior thalamus; HY, hypothalamus; SC, superior colliculus; MRN, midbrain reticular nucleus; VTA, the ventral tegmental area; SN, substantia nigra; PAG, periaqueductal gray; IC, inferior colliculus; P, pons; MY, medulla; CBN, cerebellar nuclei; CBX, cerebellar cortex; IT, intra-telencephalic; ET, extra-telencephalic.

### Distinguishability of ET-versus IT-projecting neurons across the whole brain

Are neurons in different sources that project to the same target combinations more or less distinguishable? In isocortex, the most explicit correspondence between projection types and molecular types is observed for neurons that project to ET targets versus IT targets (For a breakdown of ET and IT target regions sampled, see **Fig. 1b**.). To investigate whether such distinctions are shared with neurons from other sources, we explored the genetic distinguishability of neurons projecting to ET versus IT targets across source brain areas. To visualize the distinguishability of ET versus IT neurons from different sources we first used the joint t-SNEs of Epi-Retro-Seq and unbiased datasets of the 10 region groups (**Extended Data Figs. 2g and 4**) that contain both ET and IT neurons from the same source, and color-coded the Epi-Retro-Seq data for IT versus ET projections (**Fig. 2a**). For the cortical source t-SNE plots, the ET-projecting neurons clearly separate into a distinct cluster (L5 ET) while the IT neurons are found distributed across the annotated IT clusters, as expected. ET and IT neurons are also well-separated for the projection neurons in the entorhinal cortex (illustrated in the RHP plot) as well as for thalamic (TH) ET and IT neurons, as expected from known projections of glutamatergic TH neurons to cortex versus GABAergic neurons to subcortical targets. (See further detailed consideration of TH neurons below.) ET versus IT neurons show varying levels of separation for the other sources. While t-SNE plots allow visualization of similarities in a convenient 2-D format, they cannot fully capture the high-dimensionality of snmC-seq data. We therefore compared computed AUROC scores for ET versus IT neurons from each of the 22 sources in **Figure 2b**. Generally, comparisons show some degree of separability for each of the source regions, but AUROC scores are higher for cortical sources than for subcortical courses (except TH and AON).

**Fig 2.**
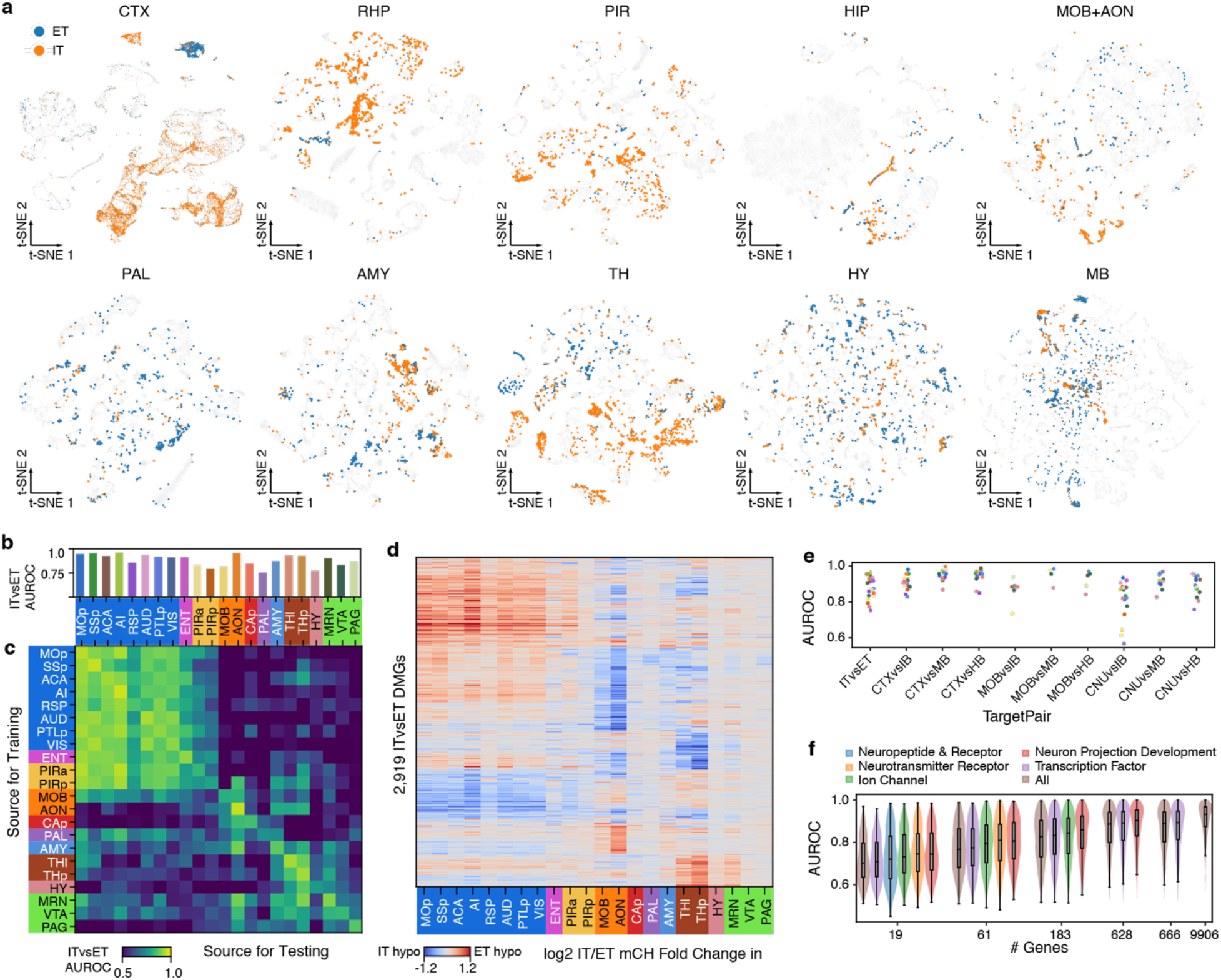
Distinguishability of neurons projecting to different targets across the entire brain. **a**, Joint t-SNEs of Epi-Retro-Seq, unbiased snmC-seq, and scRNA-seq data from 10 region groups containing both IT- and ET-projecting neurons from the same source. Only the Epi-Retro-Seq neurons projecting from the same source to ET (blue) and IT (orange) targets are colored and the other cells are in gray. RHP, retro hippocampal region. **b**, AUROC scores for comparisons between IT versus ET neurons from each of the 22 source regions. **c**, AUROC for IT versus ET neurons when the model was trained in one source region (row) and tested on another source region (column). A high AUROC indicates that the epigenetic differences between IT and ET neurons were similar between the training and testing sources. The values on the diagonal of (**c**) are the same as values in (**b**). **d**, The log2 fold changes of mCH levels in each source at the IT versus ET differentially methylated genes (DMGs). A total of 2,919 DMGs were shown that were identified in at least one of the 22 source regions. The row and column colors represent region groups in (**b-d**). **e**, AUROC for comparisons of neurons projecting to each pair of target groups. Each dot represents the comparison in one source region and the same color palette was used as in (**b**). **f**, AUROC for the comparison of all target pairs from every source region (n=926) with models using different sets of genes as features. Only subsets of the 9,906 genes with high coverage in single cells were used. The larger gene sets were downsampled to the same number of genes as the smaller sets for comparison. All comparisons between gene sets are significant (Wilcoxon signed-rank test) except the ones between “neuron projection development” and “neurotransmitter receptor” with 19 genes.

We next asked whether the epigenetic differences between ET- and IT-projecting neurons are shared across sources or alternatively whether different sources might have distinct molecular signatures that distinguish ET from IT neurons. We trained logistic regression models to distinguish ET-vs. IT-projecting neurons in each one of the 22 sources, and tested whether each model could accurately separate ET and IT neurons from each of the other sources (**Fig. 2c**). We observed that the knowledge learned by the models could largely be transferred between isocortical sources and between isocortical and archicortical (ENT and PIR) areas, but not beyond the cortical regions. Other source groups sharing similar ET vs. IT differences include MOB and AON, as well as AMY, TH and MRN. To further evaluate these relationships, we identified the differentially methylated genes (DMGs) between ET and IT-projecting cells, which merge into a combined set of 2,919 genes. Consistent with the AUROC results, these DMGs show similar fold changes across isocortical and archicortical areas, MOB and AON, as well as different parts of TH and MB (**Fig. 2d**).

The ET vs. IT differences described above group together various more specific targets, which are nevertheless distinct structures. We, therefore, assessed whether neurons projecting to more finely separated groups of targets might be more or less separable. We separated the ET and IT targets into 3 finer groups (IT: CTX, MOB, CNU; ET: IB, MB, HB) and asked which pairs of target groups are less separable between ET and IT. Most of the target group pairs have better prediction results than ET vs. IT, except that the CNU vs. IB projecting cells are less distinguishable compared to ET vs. IT based on DNA methylomes with linear models (**Fig. 2e**).

To better understand what types of genes are contributing to the predictions of projection targets, we used genes assigned to different gene ontology terms as features to compute AUROC scores. We used genes from five different categories that are considered to be associated with neuronal cell identity and projections, including: 1) neurotransmitter receptors, 2) neuropeptides and receptors, 3) ion channels, 4) transcription factors, and 5) neuron projection development (**Methods**). Since different categories have different numbers of genes, and use of more genes increases prediction performance, we downsampled the larger gene categories into samples including the same numbers of genes as the smaller categories to facilitate comparisons in 5 different groups using from 19 to 666 genes and compared the AUROC scores. We observed that the neuron projection development genes have the strongest target prediction power, followed by neurotransmitter receptors, ion channels, neuropeptides and receptors, transcription factors, and randomly selected genes (**Fig. 2f**). Using all 628 genes in the neuron projection development category achieved an average AUROC of 0.88, which is slightly lower than using all the 9906 genes as features (AUROC 0.91, **Fig. 2f**), suggesting that additional genes from other GO categories also contribute to the target predictability.

### Epi-Retro-Seq of hypothalamic projection neurons and integration with spatial assays

Although the hypothalamus is relatively small in size when compared to other profiled brain regions, analyses of gene expression and DNA methylation patterns have revealed the existence of numerous cell clusters within the hypothalamus, indicating a high level of cell type diversity (**Fig. 3a**) (See also companion papers^16,17^). Additionally, the hypothalamus is comprised of many distinct subregions and nuclei, each with unique functions and contributions to innate behaviors such as aggression, mating and feeding^18^. The hypothalamus therefore serves as an excellent use case for our data set to further examine the relationships between neuronal cell types as defined by their transcriptional and epigenomic signatures, their projection patterns, and their spatial organization.

**Fig 3.**
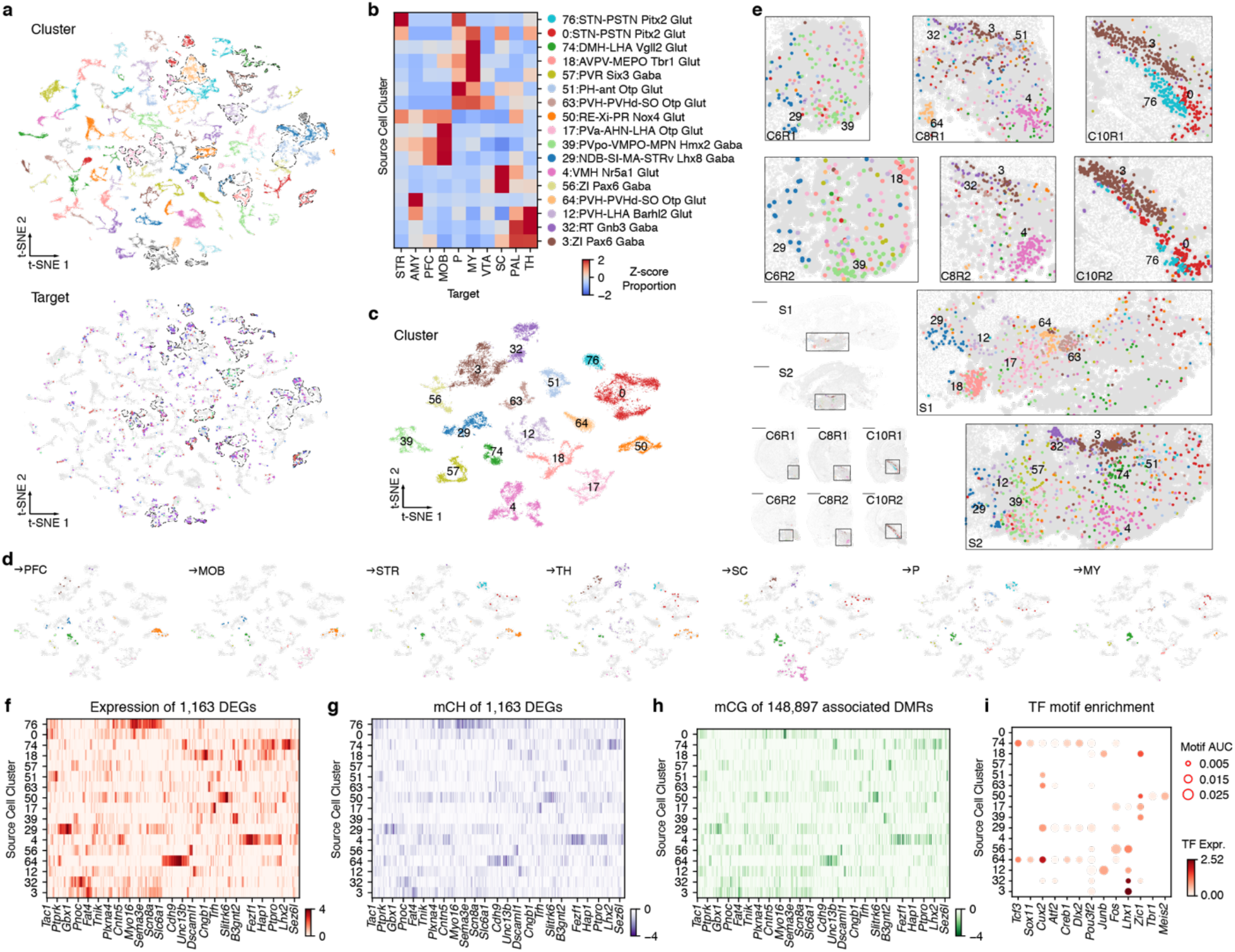
The diversity of cell type, spatial location, and gene regulation of hypothalamic projection neurons. **a**, Joint t-SNE of Epi-Retro-Seq (n=1,572), unbiased snmC-seq (n=11,554), and scRNA-seq (n=148,840) data of hypothalamic neurons colored by cell cluster (top) or projection target (bottom, same color palette as left row colors in **Fig. 1b**). Seventeen clusters enriched for the profiled projection neurons are outlined. **b**, The proportion of each of the 10 projections in each of the 17 projection-enriched clusters, Z-score normalized across targets. **c, d**, t-SNE of the 17 projection-enriched clusters (**c**), where neurons projecting to different targets were highlighted in (**d). e**, Projection-enriched HY clusters mapped to MERFISH data of 6 coronal slices (C6R1, C6R2, C8R1, C8R2, C10R1, C10R2) and 2 sagittal slices (S1, S2) of HY. The replicates of coronal slices (R1 and R2) are arranged from anterior to posterior (C6, C8, C10), left to right. The sagittal slices are arranged from lateral to medial (S1, S2), top to bottom. Examples of clusters with specific spatial locations are labeled in the enlarged insets of each slice. Scale bars represent 15 mm. The same color palette for clusters is used in (**b-e**). **f-h**, Gene expression (**f**), gene body mCH (**g**) levels of differentially expressed genes (DEGs) between the 17 projection-enriched clusters, or mCG levels of DEG-associated DMRs (**h**) in each cluster. The values are Z-score normalized across clusters. The DEGs and cell clusters are arranged in the same orders in (**f-h**). Only the DMR with highest anti-correlation with each DEG are shown in (**h**) to make the column orders consistent between (**f-h**). Examples of DEGs with GO annotations related to neuronal function and connectivity are labeled on the x-axis. **i**, Examples of transcription factors (TFs) whose binding motifs were enriched in hypo-CG-methylated DMRs are shown in the bubble plot. The size of each dot represents the enrichment level (AUC). The color of the dot indicates the expression level of the TF. The clusters are arranged in the same order as in (**f-h**).

We profiled hypothalamic neurons that project to ten distinct targets throughout the brain, including PFC, MOB, STR, PAL, AMY, TH, SC, VTA+SN (referred to later as VTA), P, and MY. By integrating Epi-Retro-Seq data with unbiased snmC-seq and single-cell RNA-seq hypothalamic data, we identified a total of 94 neuronal cell clusters, of which 17 were enriched for the profiled HY projections (**Fig. 3a,b**). Each of the projections to the ten targets was enriched in a unique subset of cell clusters, as quantified by the normalized proportion of each projection in each cluster (**Fig. 3b**). For example, HY →STR neurons were predominantly enriched in cluster 76, while HY →AMY neurons were uniquely enriched in cluster 64, indicating distinct cell type specificity of different HY projection neurons. Notably, HY neurons projecting to targets within the same major brain structure occupied overlapping yet different sets of clusters: HY →P and HY →MY were both enriched in clusters 0, 74, 18, 57, 51, and 63, but only HY →P neurons were enriched in cluster 76. Similarly, HY →PFC and HY →MOB neurons were both enriched in different subsets of cluster 50, 39, and 29, but HY →MOB neurons were uniquely enriched in cluster 17. These results suggest that HY neurons projecting to structurally related targets may share some common genetic/epigenetic cell types but also exhibit some level of diversity. In summary, HY neurons projecting to each brain region were enriched in a specific set of cell clusters (**Fig. 3b-d**). These findings underscore the cell type specificity and diversity of hypothalamic neurons projecting to different targets, shedding light on the potential functional roles of these cell clusters in various physiological and behavioral processes.

Next, we examined the spatial distributions of projection-enriched HY neuron clusters. We performed MERFISH on both sagittal and coronal brain slices to visualize the spatial location of neurons. By using the gene expression signatures of the projection-enriched clusters, we mapped them to MERFISH cells (**Fig. 3e, Methods**). Strikingly, the majority of the 17 projection-enriched clusters were located in different HY sub-regions, and the spatial distributions of cells from many clusters were distinguished by well-defined boundaries. For instance, clusters 0, 3, and 76 were located in separate “stripes” in the dorsolateral hypothalamus, in regions corresponding to Zona Incerta (ZI) or Subthalamic nucleus (STN) (**Fig. 3e**). Neurons assigned to other clusters, such as 29 and 39 (**Fig. 3e**), occupied distinct areas but were partially intermixed with neurons from other clusters. With respect to projection targets, some clusters that were enriched for particular projections were relatively confined to specific regions within the hypothalamus, while other projection-enriched clusters were distributed topographically across the hypothalamus. For example, HY →TH neurons were enriched in clusters 12, 32, and 3, all of which were located in well-delineated subregions of dorsal hypothalamus (**Fig. 3b,e**). In contrast, the seven clusters enriched for HY →P were distributed along the anterior to posterior axis of the hypothalamus and also occupied locations across the dorso-ventral and medio-lateral axes (**Fig. 3b,e**). Overall, our findings underscore the fine-scale spatial organizations of these projection-enriched cell clusters within the hypothalamus and the varying degrees of topographical heterogeneity of the locations of projection-defined HY neuronal populations.

To gain insight into the molecular characteristics and gene regulation of the projection-enriched clusters, we further utilized the integrative analysis of Epi-Retro-Seq, snmC-seq, and scRNA-seq. We identified 1,163 differentially expressed genes (DEGs) across the 17 clusters in all pairwise comparisons (**Fig. 3f**). In **Figure 3f,g**, the projection-enriched clusters are organized along the y-axis in the same order as for the target-cluster enrichment illustrations in **Figure 3b**. Each row has a largely independent pattern suggesting that each cluster has a different set of DEGs, even when there are multiple clusters enriched for projections to a particular target. (Note that this contrasts with results for TH. See below.) Notably, many of the DEGs were found to be involved in neuronal function and connectivity, as exemplified by a few highlighted genes in **Figure 3f**. As expected from the typical inverse relationship between gene expression and gene body CH methylation (mCH), mCH levels plotted with an inverted colormap in **Figure 3g** are strikingly similar to the expression levels for the same genes shown in **Fig. 3f** indicating differential methylation of these genes across clusters (**Fig. 3g**). To investigate the regulation of these DEGs, we identified 148,897 DMRs associated with the DEGs (**Methods**). The CG methylation (mCG) levels of the DEG-associated DMRs exhibited differential methylation patterns consistent with the gene expression and gene body mCH levels (**Fig. 3h**). To uncover the regulatory network of these DEGs, we further identified transcription factors (TFs) whose binding motifs were enriched in CREs (**Fig. 3i**). The analysis showed some shared sets of TFs between clusters enriched for some projections, such as HY →TH. In contrast, more varied sets of TFs were identified between clusters enriched for some other projections, such as HY →P or HY →MY. Additionally, distinct sets of TFs were observed between clusters that were enriched for different projections. Collectively, these findings underscore the existence of diverse gene regulatory networks that employ distinct TFs and DMRs for different hypothalamic projections. Furthermore, they offer valuable insights into the molecular mechanisms that govern the regulation of projection-enriched cell clusters and their associated genes in the hypothalamus.

In summary, our integrative analysis has revealed the relationships between hypothalamic neurons projecting to ten different targets and their methylation profiles, enrichment in genetic/epigenetic clusters, and spatial locations of neurons belonging to those clusters. A prior study linked transcriptomic clusters and their spatial locations within the medial pre-optic area (MPOA) of the hypothalamus to specific behaviors^19^, suggesting that those clusters might mediate their differential contributions to behavior through differences in their projections. Another study directly linked transcriptomic clusters of neurons and their locations within the ventro-medial hypothalamus (VMH) to their projections to the MPOA or PAG by combining retrograde labeling with scRNA-seq and seqFISH^6^. Those experiments revealed projection-enriched clusters, as we have found for a different set of hypothalamic projection targets, but they did not observe clear relationships between transcriptomic clusters, behavior-specific activation, and projections to the PAG or MPOA. We mapped the Kim et al. ^6^ neurons to our hypothalamus clusters and found that none of the behavior enriched clusters or projection-enriched clusters from Kim et al. correspond to any of our projection-enriched clusters in the entire hypothalamus (**Extended Date Fig. 7, Methods**). Our observations across the full spatial extent of hypothalamus and a large number of projection targets reveal strong correlations between clusters and projection targets, suggesting that cell types defined by their projections and genetics/epigenetics are also likely to make distinct contributions to hypothalamic function and related behaviors.

### Epi-Retro-Seq of thalamic projection neurons

The thalamus is a primary hub in sensory and cortical information processing and also projects to subcortical structures. Similar to hypothalamus, thalamus consists of a large number of nuclei that are organized into multiple functional groups. The main, central regions of the thalamus are composed of exclusively excitatory regions (except for a few local GABAergic interneurons in the dorsal lateral geniculate nucleus (LGd)) that are reciprocally connected with cortical areas^20^. Other more ventral and lateral regions of the thalamus (such as LGv and RT) contain GABAergic inhibitory neurons that are either reciprocally connected with thalamic excitatory neurons or project to subcortical structures such as the basal ganglia and brainstem^20^. In contrast to the hypothalamus, the thalamus had a lower degree of cell type complexity as shown by the smaller number of cell clusters identified through gene expression analysis^16^. Despite both the thalamus and hypothalamus showing a high level of heterogeneity in their anatomical nuclei and projections, the differences in their cell type complexity prompted us to investigate whether the relationships between cell types, their projections, and their spatial locations in the thalamus differ from those observed in the hypothalamus, as discussed above.

We analyzed thalamic neurons that project to twelve different targets, including nine cortical areas (PFC, MOp, SSp, ACA, AI, AUDp, RSP, PTLp, and VISp), SC, VTA, and P. To gain a comprehensive understanding of these neurons, we combined Epi-Retro-Seq data with unbiased snmC-seq and single-cell RNA-seq data from the thalamus. Through this integration, we identified a total of 58 thalamic neuronal cell clusters (**Fig. 4a**), of which 33 clusters were enriched for Epi-Retro-Seq neurons (**Fig. 4b**). It is worth noting that neurons dissected from different anatomical regions within the thalamus were located in distinct sets of clusters^17^ (**Fig. 4c**), as expected from prior descriptions based on analysis of scRNA-seq data^7^, suggesting that these molecularly defined cell clusters also have a spatial organization.

**Fig 4.**
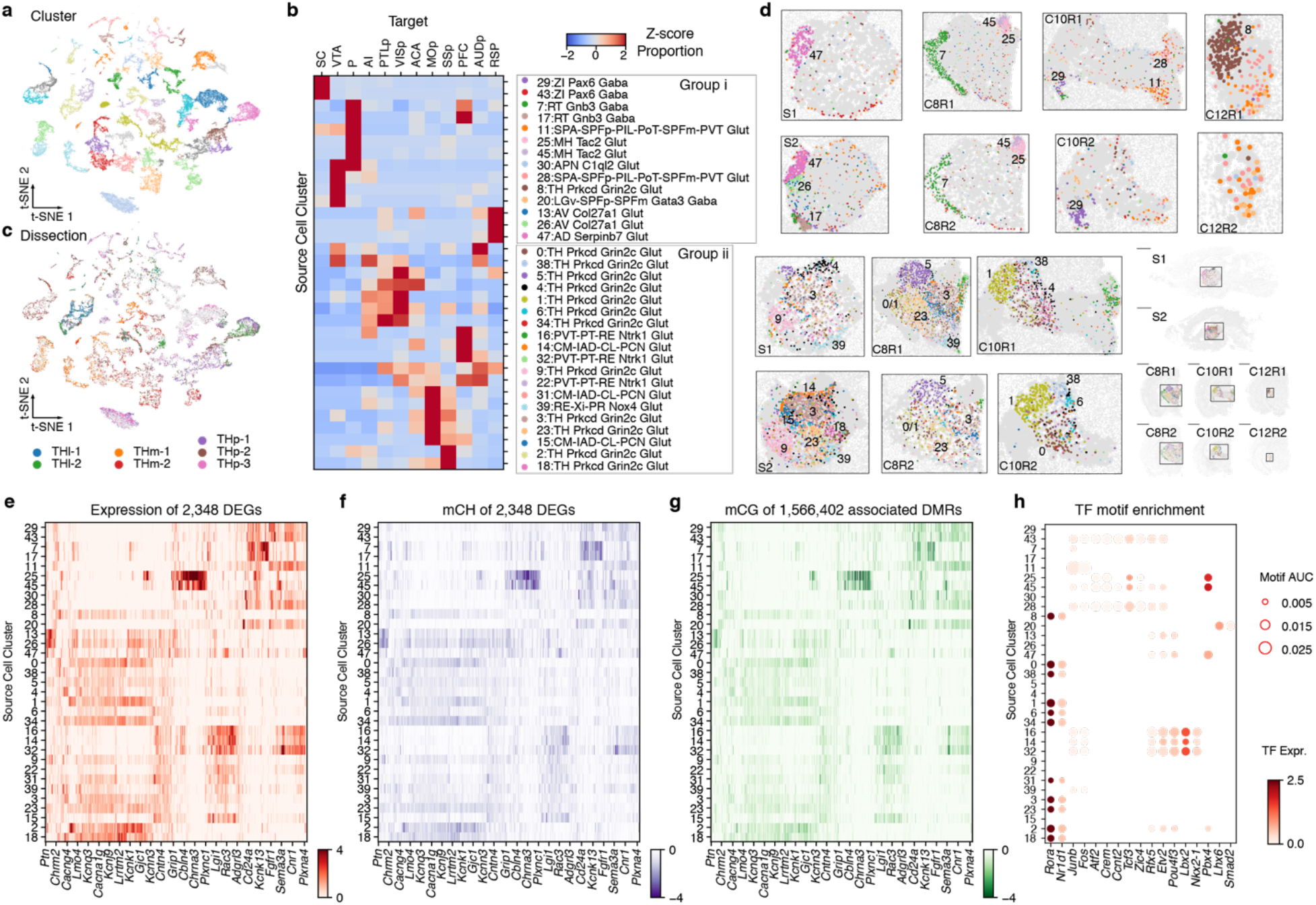
The diversity of cell type, spatial location, and gene regulation of thalamic projection neurons. **a**, Joint t-SNE of Epi-Retro-Seq (n=2,606), unbiased snmC-seq (n=16,943), and scRNA-seq (n=162,795) data of thalamic neurons colored by cell cluster. **b**, The proportion of each of the 12 assayed TH projections in each of the 33 projection-enriched clusters, Z-score normalized across targets. The same set of colors is used for labeling clusters in both cluster group i (14 clusters) and cluster group ii (19 clusters). **c**, TH neurons dissected from two anterior lateral regions (THl-1, THl-2), two anterior medial regions (THm-1, THm2), and three posterior regions (THp-1, THp-2, THp-3) are colored respectively in the t-SNE. See slices 7-10 in **Extended Data Fig. 1** for details of the dissection regions. **d**, Projection-enriched TH clusters mapped to MERFISH data of 6 coronal slices (C8R1, C8R2, C10R1, C10R2, C12R1, C12R2) and 2 sagittal slices (S1, S2) of TH. The sagittal slices are arranged from lateral to medial (S1, S2), top to bottom. The replicates of coronal slices (R1 and R2) are arranged from anterior to posterior (C8, C10, C12), left to right. The colors of clusters in the top two rows of insets are the same as the cluster labels in cluster group i in (**b**). The colors of clusters in the bottom two rows of insets are the same as the cluster labels in cluster group ii in (**b**). Examples of clusters with specific spatial locations are labeled in the enlarged insets of each slice. Note that slices C12R1 and C12R2 are not shown for cluster group ii. **e-g**, Gene expression (**e**), gene body mCH (**f**) levels of differentially expressed genes (DEGs) between the 33 projection-enriched clusters, or mCG levels of DEG-associated DMRs (**g**) in each cluster. The values are Z-score normalized across clusters. The DEGs and cell clusters are arranged in the same orders in (**e-g**). Only the DMR with highest anti-correlation with each DEG are shown in (**g**) to make the column orders consistent between (**e-g**). Examples of DEGs with GO annotations related to neuronal function and connectivity are labeled on the x-axis. **h**, Examples of transcription factors (TFs) whose binding motifs were enriched in hypo-CG-methylated DMRs are shown in the bubble plot. The size of each dot represents the enrichment level (AUC). The color of the dot indicates the expression level of the TF. The clusters are arranged in the same order as in (**e-g**).

In **Figure 4b**, we assessed the degree of enrichment of each projection in each cluster. Similar to what was observed in the hypothalamus, each population of thalamic projection neurons exhibited enrichment in distinct subsets of cell clusters, with each cluster showing enrichment for a specific set of projections, sometimes only one. Notably, the clusters enriched for TH →SC, TH →VTA, TH →P, and TH →cortex were mostly mutually exclusive. Regarding cortical projections, TH →PTLp and TH →VISp neurons exhibited enrichment in a largely overlapping set of clusters, but with varying degrees of enrichment. TH →MOp and TH →SSp neurons also shared most of their enriched clusters, which differed from those enriched for TH →PTLp and TH →VISp. These results support the notion of a separation of thalamic cell types between the visual and motor pathways in the thalamus and highlight the heterogeneity of cell types within each pathway. Notably, TH →RSP neurons showed no overlap in enriched clusters with any other cortical projections, and were uniquely enriched in clusters 13, 26 and 47. These clusters were annotated by their gene expression patterns as belonging to the anteroventral (AV) nucleus (clusters 13 and 26), and anterodorsal (AD) nucleus (cluster 47), which is consistent with TH →RSP projections originating from anterior thalamic nuclei^21^. In summary, TH neurons projecting to cortex vs. subcortical targets were enriched in distinct sets of clusters. The enriched cell clusters for cortical projections were further segregated by different thalamic pathways, with multiple enriched cell clusters observed for each pathway or projection. These findings highlight the cell type specificity as well as heterogeneity at the level of TH projections.

Such cell type specificity and heterogeneity of TH projection neurons were also reported in transcriptomic analysis of single TH projection neurons. Phillips et al. conducted a single-cell RNA-seq study on thalamic neurons projecting to the prefrontal, motor, somatosensory, auditory, and visual cortices^7^. We will refer to their dataset as Retro-Seq of these thalamocortical (TC) projections. In their study, clustering analysis of Retro-Seq revealed that neurons of each projection were enriched in a specific set of clusters, indicating specific and shared cell types between different TC projections, as well as cell type heterogeneity within each TC projection.

To compare Retro-Seq and Epi-Retro-Seq data for these TC projections, we mapped the Retro-Seq data onto our 58 integrated TH clusters (**Extended Data Fig. 8a, Methods**). The t-SNE of Retro-Seq and Epi-Retro-Seq neurons from comparisons of the most closely corresponding projections showed that they occupied similar spaces and were enriched in a common set of clusters (**Extended Data Fig. 8a,b**). However, in addition to the shared projection-enriched clusters, certain clusters were found to be enriched for each projection only in either Retro-Seq or Epi-Retro-Seq. These differences are most likely due to the use of different injection coordinates for each cortical target (**Supplementary Table 4**), resulting in overlapping yet different populations of retrogradely labeled TH neurons being analyzed in the two data sets. The degree of alignment between Epi-Retro-Seq and Retro-Seq was quantified by the overlap score and cosine distance (**Extended Data Fig. 8c, Methods**), which revealed that Retro-Seq neurons of each projection were more similar to Epi-Retro-Seq neurons for the corresponding projections than for any other projections. Taken together, these results provide further evidence for the remarkable cell type specificity and heterogeneity within each thalamocortical projection, as revealed by the analysis of both gene expression and DNA methylation.

Similar to our approach for the hypothalamus, we utilized the MERFISH data to map the spatial locations of the 33 TH projection-enriched clusters (**Fig. 4d**). Notably, almost all of these clusters exhibited a unique spatial pattern, many of them with distinct boundaries in the distributions of their cells (**Fig. 4d**). These boundaries often corresponded to specific thalamic nuclei, exemplified by clusters 25 and 45 that were enriched for pons-projecting neurons and annotated as medial habenula (MH) cell types based on their molecular signatures. When mapped to the MERFISH data, cells in these clusters demonstrated a clearly defined spatial location that corresponded to MH. This illustrates the high resolution of our data and analysis, enabling the identification of specific MH →P projection neurons among all thalamic neurons. Similarly, we were able to accurately map the molecularly annotated AD cluster 47 and AV cluster 26 that were enriched for the TH →RSP projection to their corresponding locations in the dorsal and ventral anterior thalamus. This high resolution of our data also allowed us to investigate the molecular and spatial cellular heterogeneity within a projection. For instance, the visual input from the retina reaches VISp through LGd in TH. When mapped to MERFISH, clusters 38, 5, 4, 1, and 6 that were enriched for TH →VISp neurons collectively occupied the location that corresponds to LGd, with each cluster having a unique distribution within LGd. These findings underscore the heterogeneity of LGd →VISp neurons and provide valuable insights for future in-depth analysis of different types of LGd →VISp neurons.

Next, we investigated the gene regulations of thalamic neurons in these projection-enriched clusters. Joint analysis of single-cell RNA-seq and scmC-seq data of thalamus identified a total of 2,348 differentially expressed genes (**Fig. 4e,f**) and 1,566,402 associated DMRs (**Fig. 4g**) across the 33 clusters. As expected, the expression levels of the DEGs (**Fig. 4e**) were anti-correlated with their mCH levels (**Fig. 4f**), while their associated DMRs also showed strong correspondence in terms of mCG levels (**Fig. 4g**). In contrast to HY, TH clusters enriched for the same projections displayed similar expression patterns of DEGs and methylation patterns of the associated DMRs, suggesting shared usage of CREs in gene regulation. Additionally, we identified transcription factors with significant motif enrichment in the DMRs (**Fig. 4h**). Clusters enriched for the same projection had similar sets of TFs, while those enriched for different projections had more distinct sets of TFs. These results imply the existence of projection-specific gene regulatory networks, which consist of unique sets of TFs, CREs, and target genes in the thalamus. These relationships are in contrast to those observed in HY, where the organization of TF motifs is not closely related to projection targets.

### Differential usage of neurotransmitters in projection neurons

Recent brain-wide single-cell and spatial transcriptomic analyses have revealed remarkable heterogeneity and spatial specificity in neurotransmitter usage among different cell types across the mouse brain^16,17^. As described above and exemplified in thalamus and hypothalamus, our integrative analysis revealed high levels of cell-type and spatial specificity in neurons with different projections. These findings sparked a further investigation into the neurotransmitter usage of these distinct projection neurons that were in different brain regions and had different cell type compositions. Insights into the neurotransmitter usage of different projection neurons may shed light on their functional properties and their potential role in behavior, with broader implications for understanding neural circuits and the mechanisms underlying various brain functions and disorders.

To systematically examine the use of neurotransmitters by different projections, we quantified the levels of expression of nine canonical neurotransmitter transporter genes in each of the projection-enriched clusters within the twelve grouped brain regions described previously (**Extended Data Fig. 5**). These transporter genes included *Slc17a7* (*Vglut1*), *Slc17a6* (*Vglut2*), and *Slc17a8* (*Vglut3*) for glutamatergic neurons, *Slc32a1* (*Vgat*) for GABAergic neurons, *Slc6a2* (*Net*) for noradrenergic neurons, *Slc6a3* (*Dat)* for dopaminergic neurons, *Slc6a4* (*Sert*) for serotonergic neurons, *Slc6a5* (*Glyt2*) for glycinergic neurons, and *Slc18a3* (*Vacht*) for cholinergic neurons. In addition, we used histidine decarboxylase (*Hdc*) for histaminergic neurons. Our analysis revealed a diverse range of neurotransmitter usage across the projection-enriched clusters, particularly those in the midbrain and hindbrain regions. Furthermore, a large proportion of the projection-enriched clusters exhibited significant expression of more than one neurotransmitter transporter gene. These findings support that there is a wide variation in neurotransmitter usage across different neural pathways and highlight the heterogeneity within some of these pathways. Below, we delve deeper into a few interesting cases, including projections from the hindbrain regions of P and MY, the amygdala, and the midbrain region of VTA.

#### Epi-Retro-Seq and Neurotransmitter Usage in Hindbrain Projection Neurons

We analyzed eleven hindbrain projections, which included projections from P or MY to five different targets -TH, HY, SC, CBN, and CBX - as well as the projection from P to MY. These projections were enriched in 20 cell clusters out of a total of 128 hindbrain clusters. The degree of enrichment of each projection in each cluster was quantified as shown in **Figure 5a**. Notably, in both P and MY, neurons projecting to the CBX were the most distinct from other projection neurons. This was evidenced by the presence of exclusive CBX-projecting clusters in each region. For instance, in P, the cluster 0 was uniquely enriched for the P**→**CBX projection, while in medulla, cluster 76 was enriched for MY neurons projecting to the cerebellum, particularly those projecting to CBX.

**Fig 5.**
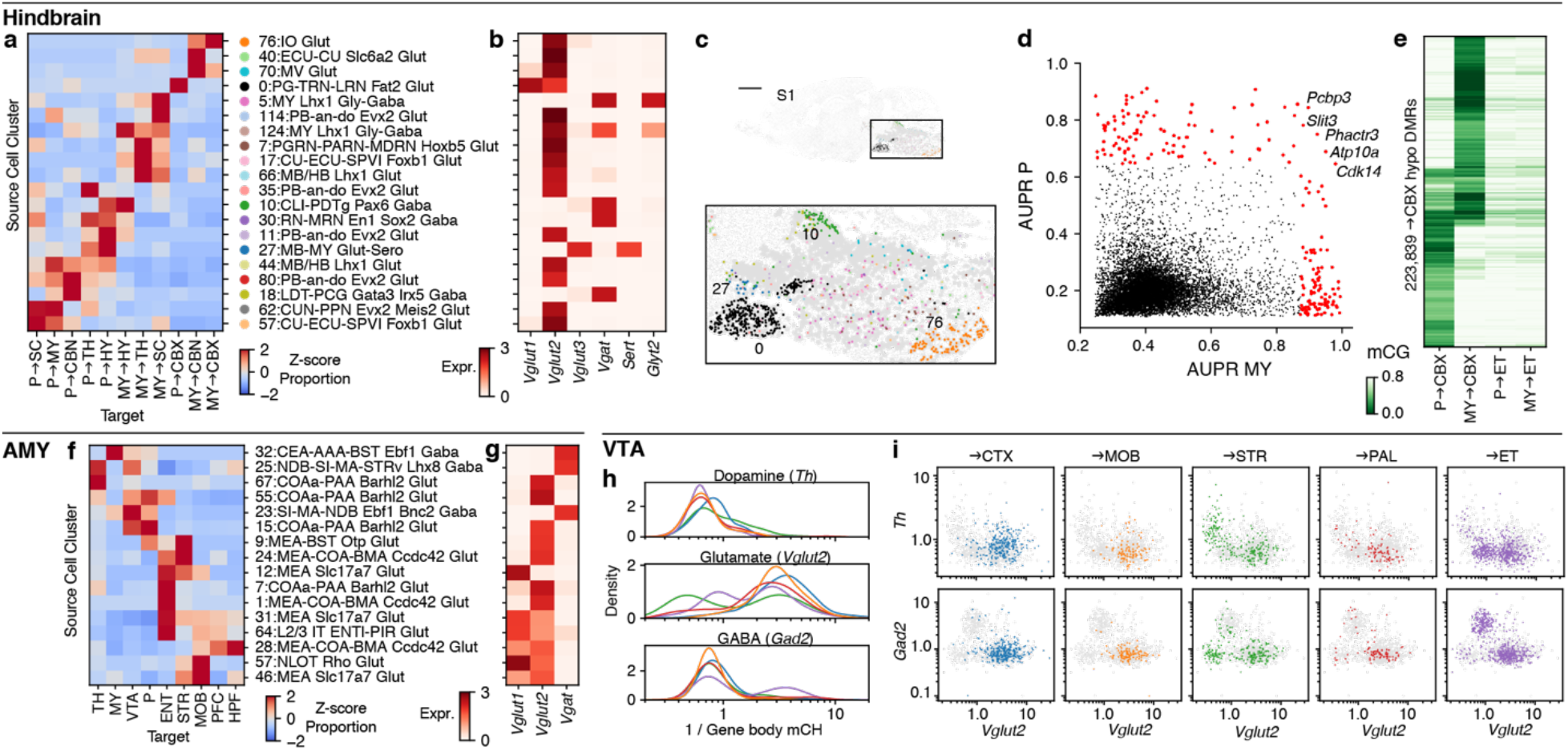
Neurotransmitter usage in HB, AMY, and VTA projection neurons. **a, b**, The proportion of each of the 11 assayed HB projections (**a**, Z-score normalized across targets) and the expression levels of six neurotransmitter transporter genes (**b**) in each of the 20 hindbrain projection-enriched clusters. **c**, Projection-enriched clusters mapped to the MERFISH slice S1. The same color palette for clusters is used in **a** and **c. d**, Area under the curve of precision-recall (AUPR) of genes to distinguish P →CBX cluster (0) and P →ET clusters (10, 11, 27, 30, 35, 44, 57, 62, 80) vs. MY →CBX cluster (76) and MY →ET clusters (5, 7, 10, 11, 17, 57, 66, 114) with gene body mCH level in Epi-Retro-Seq data. The genes with AUPR>0.872 in P and AUPR>0.647 in MY (>99th quantile) are colored in red. The name of five genes selected in both P and MY are labeled. **e**, mCG levels of hypo-mCG DMRs in P and MY between the →CBX clusters and →ET clusters. **f, g**, The proportion of each of the 9 assayed AMY projections (**f**, Z-score normalized across targets) and the expression levels of neurotransmitter transporters *Vglut1, Vglut2*, and *Vgat* (**g**) in each of the 16 projection-enriched clusters. **h, i**, The gene-body mCH levels of tyrosine hydroxylase (*Th*), *Gad2*, and *Vglut2* in VTA projection neurons, shown in density plots (**h**) or scatter plots (**i**). Colors represent VTA neurons projecting to different targets and the same palette is used in (**h, i**). Note that, because low mCH levels indicate high gene expression, the x-axis in **h** and both axes in **i** are plotted as the reciprocal mCH values (1/gene body mCH), so low mCH is plotted to the right/up and high to the left/down. ACA was not included in CTX (see **Extended Data Fig. 9**).

The 20 projection-enriched clusters showed expression of six neurotransmitter transporter genes (**Fig. 5b**). The majority of these clusters contain glutamatergic neurons expressing *Vglut2*, such as the previously mentioned MY**→**CBX enriched cluster 76. Interestingly, *Vglut1* and *Vglut2* were co-expressed in cluster 0 neurons that were enriched for the P**→**CBX projection. These observations are consistent with previous studies that demonstrated the presence of VGLUT1 or VGLUT2 in climbing fiber (MY**→**CBX) terminals and both VGLUT1 and VGLUT2 in cerebellar mossy fiber (P →CBX) terminals using synaptic vesicle immunoisolation^22^. Moreover, different neurotransmitters were utilized in clusters enriched for the same projections. For instance, clusters 10, 30, 11, and 27 were enriched for P**→**HY projections. Among them, clusters 10 and 30 are GABAergic with *Vgat* expression, cluster 11 is glutamatergic with *Vglut2* expression, whereas cluster 27 is serotonergic showing co-expression of *Sert* and *Vglut3*. Furthermore, several of these clusters also exhibited distinctive spatial distributions when mapped to the MERFISH data, such as clusters 0, 76, 10, and 27 (**Fig. 5c**). Altogether, these results underscore the extent of molecular, cellular, and spatial specificity and diversity within hindbrain projections.

We observed that neurons projecting to CBX from P or MY were distinct from other projections originating from the same regions. To investigate this further, we examined the molecular signatures that could differentiate CBX-projecting neurons from other projection neurons in P or MY. Additionally, we investigated whether there were any common molecular signatures between the P**→**CBX and MY**→**CBX projections. Analysis of gene body DNA methylation identified genes that could distinguish the P**→**CBX cluster (0) from other P projections associated clusters, or differentiate MY**→**CBX cluster (76) from other MY projections associated clusters (**Fig. 5d**). Interestingly, only five genes were common between the top 100 genes in the two sets, namely *Slit3, Phactr3, Pcbp3, Atp10a*, and *Cdk14* (highlighted in **Fig. 5d**). Notably, *Slit3* encodes a repulsive axon guidance molecule^23,24^, and *Phactr3* has been shown to be involved in regulating axonal morphology^25,26^. To understand how the differentially expressed genes in CBX-projecting neurons are regulated, we explored DMRs that were hypo-methylated in CBX-projecting neurons. In total, we identified 223,839 hypo-DMRs in the hindbrain that were associated with CBX-projecting neurons (**Fig. 5e**). These DMRs were further divided into subsets that were hypo-methylated in either P**→**CBX or MY**→**CBX, while only a limited number were hypo-methylated in both. Collectively, these findings suggest that the molecular mechanisms underlying CBX versus other projections in P and MY are largely distinct, but with some shared features at both the transcriptomic and epigenomic levels.

#### Epi-Retro-Seq and Neurotransmitter Usage in Amygdala and Midbrain Projection Neurons

We examined projections from the amygdala to nine different targets, including the PFC, ENT, HIP, MOB, STR, TH, VTA, P, and MY. These projections were enriched in 16 amygdala clusters, with distinct sets of clusters enriched for neurons projecting to IT targets (ENT, STR, MOB, PFC, and HIP) versus ET targets (TH, MY, VTA, and P) (**Fig. 5f**). The clusters enriched for IT projections were primarily glutamatergic and expressed *Vglut1* and/or *Vglut2*(**Fig. 5g**). In contrast, the clusters enriched for ET projections were divided between glutamatergic clusters that expressed *Vglut2* and GABAergic clusters (**Fig. 5g**). Notably, the AMY →ENT projection was particularly distinct compared to other IT projections, with unique enrichment in clusters 1 and 7 (**Fig. 5f**). Additionally, it exhibited varied usage of vesicular glutamate transporters. Within the clusters enriched for AMY →ENT, *Vglut1* was predominantly expressed in cluster 12, *Vglut2* was the predominant transporter in clusters 24, 7, and 1, while Clusters 31 and 64 expressed both *Vglut1* and *Vglut2*, suggesting a potential diversity in the physiology and function of amygdala neurons projecting to the entorhinal cortex. In summary, our results underscore the heterogeneity in neurotransmitters and their transporter utilization among amygdala projection neurons.

The midbrain regions containing the ventral tegmental area (VTA) and substantia nigra (SN) (which we collectively refer to as VTA) exhibit some of the most interesting and complex patterns of heterogeneous neurotransmitter usage between different projections. Our study analyzed VTA neurons projecting to sixteen different targets, including six cortical targets (PFC, MOp, SSp, ACA, RSP, and PTLp), six other IT targets (MOB, ENT, PIR, AMY, STR, and PAL), and four ET targets (TH, HY, SC, and P). By integrating Epi-Retro-Seq and unbiased snmC-seq data, as well as single-cell RNA sequencing of VTA, we can distinguish between cell clusters with various combinations of the expected glutamate, GABA, and dopamine transporters known to be expressed by VTA neurons^27–30^ (**Extended Data Fig. 9a,b**).

In order to better examine the relationships between VTA neurons projecting to different targets and their use of neurotransmitters, we analyzed the levels of mCH at specific marker genes, including tyrosine hydroxylase (*Th*) for dopaminergic neurons, *Gad2* for GABAergic neurons, and *Vglut2* for glutamate neurons because previous studies showed that rodent VTA glutamate neurons mainly express *Vglut2* but not *Vglut1* or *Vglut3*^*31,32*^ (**Fig. 5h,i and Extended Data Fig. 9c-d**). Lower mCH levels at these genes suggest higher gene expression, given the negative correlation between gene body mCH levels and gene expression. In general, VTA neurons that project to the cortex had lower levels of mCH at *Th* compared to subcortical projections (except for VTA →STR), suggesting a higher expression of *Th* (**Fig. 5h top**, *P* values=2.8e-7 (CTX vs MOB), 3.0e-5 (CTX vs PAL), 6.2e-15 (CTX vs ET), Wilcoxon rank-sum tests). The CTX-projecting neurons also exhibited lower mCH levels at *Vglut2*, indicating a significant *Vglut2* expression (**Fig. 5h middle**).

Therefore, these CTX-projecting VTA neurons are likely *Th*+ and *Vglut2*+ and utilize both dopamine and glutamate (**Fig. 5i and Extended Data Fig. 9c**). In contrast, VTA neurons projecting to STR do not appear to co-express *Th* and *Vglut2* but instead were shown to consist of three subpopulations: *Th*+ *Vglut2*-, *Th*-*Vglut2*+, and *Th*-*Gad2*+, as indicated by their mCH levels at *Th, Vglut2*, and *Gad2* (**Fig. 5i and Extended Data Fig. 9c-e**). Based on their mCH levels, the ET-projecting neurons were generally divided into two subgroups: *Gad2*+ and *Vglut2*+ (**Fig. 5i**).

Among the ET-projecting VTA neurons, those projecting to TH and HY were more similar to each other than to those projecting to SC and P (**Extended Data Fig. 9b**). Notably, some of the SC- and P-projecting neurons were uniquely present in a VTA *Gad2*+ cluster that were absent in other projections (**Extended Data Fig. 9b**). Overall, our findings corroborate prior reports of diverse populations of VTA neurons that employ single or combined neurotransmitters and highlight intricate patterns of distinct neurotransmitter usage among various projections.

## Summary

Altogether, we have uploaded and made available data that informs potential users about the relationships between axonal projection status and DNA methylation at single-cell resolution for tens of thousands of neurons corresponding to hundreds of source →target combinations. We have provided quantitative measures of the discriminability of source neurons projecting to different targets for nearly one-thousand target-to-target comparisons. We have further demonstrated how these data can be integrated with other single-cell data modalities, including scRNA-seq and MERFISH, to link the projection status of spatially-resolved cell-type clusters to neural circuits. More extensive details about the use of these data, their potential limitations, and the analytic approaches we have taken can be found in **Methods**. The in-depth analyses provided here for both brain-wide comparisons of ET-versus IT-projecting neurons, and for the full sets of targets assayed for 6 of the 32 assayed source regions (HY, TH, P, MY, AMY, VTA) exemplify the utility of the much larger data set for further brain-wide and source-or target-focused analyses.

## Methods

### Experimental Animals

As described by Zhang et al. ^5^, all experimental procedures using live animals were approved by the Salk Institute Animal Care and Use Committee. The knock-in mouse line, R26R-CAG-loxp-stop-loxp-Sun1-sfGFP-Myc (INTACT) used in Epi-Retro-Seq^5^ was maintained on a C57BL/6J background. 42-49 day old adult male and female INTACT mice were used for the retrograde labeling experiments. Adult C57BL/6J “wild-type” mice were used for MERFISH experiments.

### Surgical Procedures for Viral Vector and Tracer Injections

As described by Zhang et al.^5^, to label neurons projecting to regions of interest, injections of rAAV2-retro-Cre (produced by Salk Vector Core or Vigene, 2×10^12^ to 1×10^13^ viral genomes/ml, produced with capsid from Addgene plasmid #81070 packaging pAAV-EF1a-Cre from Addgene plasmid #55636) were made into both hemispheres of the INTACT mice. In summary, animals were anesthetized with either ketamine/xylazine or isoflurane and placed in a stereotaxic frame. Pressure injections of 0.05 to 0.4 microliters of AAV per injection site were made using glass micropipettes (tip diameters ∼10-30μm) targeted to stereotaxic coordinates corresponding to MOp, SSp, ACA, AUDp, RSP, PTLp, VISp, HPF, MOB, STR, PAL, TH, SC, VTA+SN, P, MY, and CBX. To precisely target PFC, AI, ENT, PIR, AMY, HY, and CBN, AAV was injected using iontophoresis to ensure confined viral infection. Iontophoretic injections (+5μ;A, 7 s on/7 s off cycles for 5-10 min) were made with glass pipettes with tip diameter of ∼10μm. For most of the desired target areas, injections were made at different depths, and/or at different AP or ML coordinates to label neurons throughout the target area. More detailed injection coordinates and conditions are listed in **Supplementary Table 1**. At least 2 male and 2 female mice were injected for each desired target.

### Brain dissection

Brain dissections were done as described in Zhang et al.^5^. In summary, approximately two weeks after the AAVretro injection, brains were extracted from the 56-63 day old INTACT mice, immediately submerged in ice-cold slicing buffer (2.5mM KCl, 0.5mM CaCl_2_, 7mM MgCl_2_, 1.25mM NaH_2_PO4, 110mM sucrose, 10mM glucose, and 25mM NaHCO_3_) that was bubbled with carbogen, and sliced into 0.6 mm coronal sections starting from the frontal pole. From each AAVretro-injected brain, the slices were kept in the ice-cold dissection buffer from which selected brain regions (**Fig. 1b**) were manually dissected under a fluorescent dissecting microscope (Olympus SZX16), following the Allen Mouse Common Coordinate Framework (CCF), Reference Atlas, Version 3 (2015) (**Extended Data Fig. 1**). The dissected brain tissues were transferred to prelabeled microcentrifuge tubes, immediately frozen in dry ice, and subsequently stored at -80°C.

### Nuclei preparation and single-nucleus isolation

Nuclei preparation and isolation were done as described by Zhang et al.^5^. In summary, for each dissected brain region, samples from 2 males and 2 females were pooled separately as biological replicates for nuclei preparation. Nuclei were prepared using a modified protocol as reported by Lacar et al., 2016^33^ and described by Zhang et al.^5^. Nuclei suspensions were then incubated with GFP antibody, Alexa Fluor 488 (Invitrogen, A-21311), and anti-NeuN antibody (EMD Millipore MAB377) conjugated with Alexa Fluor 647 (Invitrogen A20173). GFP^+^/NeuN^+^ single nuclei were isolated using fluorescence-activated nuclei sorting (FANS) on a BD Influx sorter or a BD Aria Fusion cell sorter with 100μm nozzle, and sorted into 384-well plates with digestion buffer for snmC-seq. The collected plates were incubated at 50°C for 20 minutes and then stored at -20 °C.

### snmC-Seq library preparation

The bisulfite conversion and library preparation were performed following the detailed snmC-seq protocol as previously described^14^. In brief, DNA samples from single nuclei were barcoded with random primers after the bisulfite conversion, pooled through two rounds of cleaning up with SPRI beads, then added with adapters and PCR amplified to generate the libraries. Libraries were then pooled, cleaned up with SPRI beads, normalized and sequenced on Illumina Novaseq 6000 using the S4 flow cell 2 × 150 bp mode.

### Mapping and Preprocessing

Epi-Retro-Seq data were mapped to the mm10 genome as described in our previous study^34^. For each single cell, we counted the methylated and total basecalls for all 100kb non-overlapping genomic bins and all gene bodies expanded 2kb in both directions using ALLCools generate-dataset. The data is saved in Zarr format to allow chunk loading and on-disk computing^35^. To avoid the methylation differences being driven by the active and inactive X-chromosomes, we only used the autosomal bins and genes in our analyses. The cell-by-bin and cell-by-gene posterior methylation levels were computed as previously described^34^, which is the input for all downstream analyses.

### Quality control

Step 1. The cells included in the analysis are required to have 1) median mCCC level of the experiment <0.025, 2) 500,000 < nonclonal reads < 10,000,000, 3) mCCC level <0.05. In total, 56,843 cells from 703 experiments satisfied these requirements (**Extended Data Fig. 2a, b**).

Step 2. The potential doublets were removed as described in the next section, and 48,032 cells remained in the dataset (**Extended Data Fig. 2c, d**). The cell type and dissection information of these cells were used in our analysis, but further filters were applied to exclude non-neuronal cells as well as neurons whose projection targets are not confidently assigned.

Step 3. The experiments with less than 20 neurons were excluded to ensure the statistical power of projection analysis, resulting in 39,461 cells from 519 experiments left. The non-neuronal cells are also removed from the dataset, after which 34,643 neurons remain. The cell type classification method is described in the next section.

Step 4. The cortical cells from 286 experiments were further filtered to exclude the experiments with a high proportion of neurons of the cell types known not to project to the intended injection site (off-target clusters), using the same method as in our previous study^5^. Specifically, for each FANS run, we counted the number of neurons that were observed in known on-target cell types (o_on_) and off-target cell types (o_0ff_). Assuming that the proportions of contaminated cells in each subclass would be similar to a sample without projection-type enrichment, we compared the observed counts to the counts from unbiased snmC-seq data (*E*_on_ and *E*_off_) collected from the corresponding dissections in **Extended**

**Data Fig. 1**. The fold-enrichment was computed as 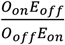 A one-sided exact binomial test of goodness-of-fit was used to determine whether the enrichment of on-target cells was significant. The *P* value was computed as *Pr*(*X* ≥ *O*_on_; *n, p*), where 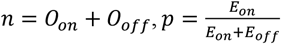. For each ET target, we considered ET as on-target subclass and IT+inhibitory neurons as off-target. The thresholds for fold-enrichment and FDR (Benjamini-Hochberg procedure) were 8 and 0.001. For IT targets, we considered IT as on-target subclasses and L6 CT+inhibitory neurons as off-target. The thresholds for fold-enrichment and FDR (Benjamini-Hochberg procedure) were 3 and 0.001. This eliminated 32 out of 286 sorting cases (**Extended Data Fig. 2e**).

The rationale of step 4 is to remove potential contamination in the dataset that might have resulted from 1) inaccurate gating of GFP+ NeuN+ cells, and 2) AAV-retro injection pipettes that passed through overlying source brain regions and directly labeled neurons at those sources rather than being taken up retrogradely from the intended target. 1) could be more common in the experiments of some weak projections, where very few neurons were retrogradely labeled, resulting in small proportions of cells passing FANS gating criteria and subsequent inclusion of high proportions of cells accepted from the edges of FANS gates. 2) could be more common when targeting a deep structure in the brain (e.g. TH, HY) and collecting cells from the superficial structures directly above the target (e.g. cortex). Note that step 4 was only performed on experiments of isocortical neurons, given that the on-target and off-target clusters were relatively clear in these areas. For subcortical projections, comprehensive prior knowledge of molecular cell types associated with projection is usually lacking, which makes the estimation of contamination using this method more challenging. The projections profiled in the subcortical structures are usually strong and do not involve overlaying of sources and targets, which would potentially lead to lower noise level in those data. Nevertheless, it is worth noting that even after these QC steps, there are still expected to be some contaminated cells remaining in the dataset. After all the QC steps, 33,304 neurons from 487 experiments were used for analyses related to projection targets.

### Transfer of cell labels from one dataset to another with weighted k-nearest neighbors

This method is similar to the label transfer method in Seurat v3^36^, and implemented in our ALLCools python package. This is used in multiple analyses throughout the manuscript, including Epi-Retro-Seq cell classification and doublet removal, and mapping of MERFISH cells and Retro-Seq cells into major dissection regions or RNA and mC co-clusters. The original Seurat method identified anchors between two datasets, and used the 100 nearest anchors for each cell in the unlabeled dataset to average the information from the labeled dataset. Since the 100 anchors usually include cells from other clusters, especially for a cell in an underrepresented cluster, this method makes the label transfer of small clusters quite noisy. Instead of using the anchors between datasets to transfer the labels, we only used the anchors to integrate the datasets together, and directly find the neighboring cells of the unlabeled dataset in the labeled dataset on the integrated space. Since the larger dataset usually has more cells than the number of anchors, this method reduced the noise in the small clusters.

Assume we have two datasets in a coembedding space, A with labels and B without labels. For each cell in B as a query cell, we first find its k nearest neighbors in A with Euclidean distance, and denote its distances to the neighbors as a k-dimensional vector *D*. *D* is then transformed to *W* as the weights for averaging the information from the neighbors through the following steps which are the same as in Seurat. 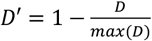; 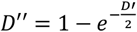; 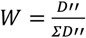 After the transformation, the closer neighbors have higher weights, and the weights of all neighbors sum up to 1. To transfer a categorical label from A to B, we used one-hot encoding to represent the label and the label vectors corresponding to the k neighbors in A of the query cell (k-by-#categories, denoted as *L*_*ref*_) were averaged with the weights *W*. The resulting vector *L*_*qry*_ *= WL*_*ref*_ represents the probability of the query cell belonging to each category. The category with the maximum probability is used as the final assignment.

### Cell classification and doublet removal

As described in our companion manuscript, the cell clustering of the unbiased dataset was performed iteratively at four levels (L1-L4), which assigned the cells into 61 (L1), 411 (L2), 1346 (L3), and 2573 (L4) clusters, respectively. At each level, the highly variable 100kb bins were selected, and PCA was used for dimension reduction. The significant PCs from mCH and mCG were combined to perform consensus clustering. We first performed doublet removal with the help of unbiased data. The 56,843 cells after QC step 2 are mapped to the 310,605 unbiased snmC-seq cells (including predicted doublet cells). We used the highly variable features selected in the unbiased data and the PCA model fit with the unbiased data to transform the Epi-Retro-Seq to the same dimension reduction space as the unbiased data. Then we classified the Epi-Retro-Seq cells into either one of the 61 L1 clusters or the predicted doublet clusters defined in the unbiased data. The classification was performed with the k-nearest neighbor approach described above on the PCs combining mCH and mCG. The Epi-Retro-Seq cells assigned to each non-doublet L1 cluster were analyzed in the next iteration, using the highly variable features selected in the unbiased data of the cluster and the PCA model fit with the unbiased data of the cluster. All the predicted doublet cells in the unbiased data were added in each L1 cluster in the level 2 clustering to further exclude the potential doublets. After these two iterations, the cells predicted to be doublets were removed, with 48,032 Epi-Retro-Seq cells remaining. These cells were mapped to the 301,626 unbiased snmC-seq cells (without predicted doublets) with the same feature selection and PCA methods through the four levels, so each Epi-Retro-Seq cell is assigned to one cluster at each level. The 61 L1 clusters were annotated based on their dissection source and marker genes. The cell clusters representing non-neuronal cells were removed from further analyses. The cells corresponding to the IT, ET, CT, and cortical inhibitory clusters in the L1 cluster annotation were used for QC step 4 as described above.

### Quantification of projection neuron difference with AUROC

To test the similarity of two groups of cells based on DNA methylation, we trained logistic regression models to predict the group label of each cell. We compared the results using four different types of features to predict the projection target of neurons from the same source. These include the posterior of 100 kb-bin mCH level, gene body mCH level, and the dimension reduction results of the two matrices. 50 PCs were used as dimension reduction, with unbiased snmC-seq to fit the PCA models and transform the Epi-Retro-Seq data. We also used two methods to split the cells into training and testing sets. One used a random selection of half of the cells projecting to each target for training and the other half for testing (computational replicates), the other was based on the sex of the mice where the cells were collected (biological replicates). After the QC steps, we have 168 source-target combinations with data from both sexes and the other 57 with cells only from one sex. Therefore, all the comparisons of 926 target pairs could be quantified with the computational replicates, but only 516 of them could be quantified with biological replicates. We noticed significant congruence of model performance between the different features and different train/test splits (**Extended Data Fig. 3a-c**). Using 100 kb-bins performed very similar to gene bodies (**Extended Data Fig. 3a**). Using raw features performed slightly better than using principal components (**Extended Data Fig. 3b**). Using computational replicates performed significantly better than biological replicates (**Extended Data Fig. 3c**), which was expected given that the computational replicates dismissed the heterogeneity between biological replicates and made the predictions easier. Nevertheless, the computational replicates still provided strongly correlated results to biological replicates (**Extended Data Fig. 3c**), which allowed the comparison between different target pairs to evaluate their epigenomic differences.

All the other results in the figures were computed using the computational replicates with gene-body mCH as features. The features were filtered based on average read coverage across cells before the model training. We removed the 100 kb bins and genes with <500 average CH basecalls, resulting in 23,730 bins or 9,906 genes in the model. Sci-kit learn was used for model implementation. The area under the receiver operating characteristic (AUROC) from cross-validation was used to measure the performance of the model. The higher AUROC represented the better ability of the model to present the group label, which indicated the two groups had larger mCH differences and were more distinguishable. For computational replicates, we performed random sampling 50 times with different seeds, and used the average AUROC as the final result.

To test the predictability of projection targets with genes from different categories, we collected the genes from the following resources. Neuropeptide and receptors: Table 1 in Smith et al.^37^ and Supplementary Figure 16 in Tasic et al^38^. Neurotransmitter receptors: Supplementary Figure 15 in Tasic et al^38^. Ion channels: Supplementary Figure 14 in Tasic et al^38^. and the Guide to PHARMACOLOGY database (https://www.guidetopharmacology.org/DATA/targets_and_families.csv). Neural projection development: gene ontology terms GO0031175 Neuron Projection Development and GO0050808 Synapse Organization. Transcription factors: annotation from SCENIC+^39^. Only genes included in 9,906 genes with high CH coverage were analyzed, and adding more lower coverage genes to increase the size of genesets did not improve the prediction performance.

Several reasons could contribute to a low prediction performance. Biological reasons would include: 1) Some neurons make projections to several targets simultaneously. These could result in the neurons being captured by multiple retrograde labeling experiments of different targets. It would be impossible to predict a single label with our pairwise models for this type of neuron. 2) Some neurons project to different target regions but have tiny epigenetic differences. To systematically distinguish 1) to 2), other anatomic and genetic validations are still needed.

Technical reasons would include: 1) The contamination levels of some experiments might be relatively high, which make larger noise and hinder the models from capturing real projection differences. 2) The epigenetic differences between neurons projecting to different targets varies across replicates. 3) The sample sizes of some projections are small, which makes learning more challenging. 4) The models are not powerful enough to capture the complex differences between projections.

Elimination of contaminated FANS runs in QC step 4 decreased the potential influence by 1) for cortical neurons as discussed in the QC section, although there are still contaminated cells included in the dataset. The improvement in labeling efficiency and specificity would help to better solve the molecular differences between projection types. In this study, male and female mice were treated as biological replicates after removing sex chromosomes. Although methylation patterns of autosomes are similar, differences between sexes or animals might still exist. The small differences in performances between data splitting methods (based on computation or biological replicates) might suggest a less notable effect contributed by 2) in those samples. To evaluate the potential limitation of 4), more carefully curated models, and accordingly, more samples, would be required. Thus, given all these factors, we are generally more confident in the distinguishable target pairs when training and testing sets were split based on both computational and biological replicates. The interpretation of comparisons without biological replicates and the indistinguishable pairs would need to be more careful and are not involved in the major conclusions in this manuscript. Our study aims to provide a general view across multiple sources and targets. A more detailed understanding of specific projections would require larger scale profiles on those specific projection types.

### Integration between snmC-seq, Epi-Retro-Seq, and scRNA-seq

snmC-seq and scRNA-seq are comprehensive atlases of the whole mouse brain, so most of the cell types are expected to be presented in both datasets. Therefore, the two datasets were integrated based on a canonical correlation analysis (CCA) framework, which captures the shared variation between the two datasets^36^. Epi-Retro-Seq is a projection-enriched dataset that contains part of the cell types in the atlas, but the shared methylation modality with snmC-seq allowed it to be integrated with the comprehensive atlas with a reciprocal PCA framework. Both the Epi-Retro-Seq and the scRNA-seq datasets were mapped to the dimension reduction space of the snmC-seq data to create a multi-modality atlas of each brain region group.

For each region group, we selected cells from the three datasets belonging to the dissection regions. The methylation cells in the L1 clusters corresponding to cerebellar neurons were excluded from the analysis of cerebral and brainstem regions. The RNA cells from the major classes of non-neuronal cells and immature neurons, and the subclasses of cerebellar neurons were excluded from the analyses. The RNA cells from subclasses of MM and DCO were also excluded due to the dissection differences between the two studies.

The gene expression levels of scRNA-seq cells were normalized by dividing the total UMI count of the cell and multiplying the average total UMI count of all cells, and then log-transformed. The posterior gene-body mCH level of snmC-seq and Epi-Retro-Seq cells were used. The cluster-enriched genes (CEGs) were identified in each L4 cluster. We checked the variance of the mCH CEGs among the snmC-seq cells and scRNA-seq cells and only used the CEGs with mCH variance greater than 0.05 and expression variation greater than 0.005 for the analyses. The opposite of mCH levels was used for snmC-seq and Epi-Retro-seq data due to the negative correlation between gene body DNA methylation and gene expression. We fit a PCA model with the snmC-seq cells and transformed the Epi-Retro-Seq cells and scRNA-seq cells with the model. The PCs were normalized by the singular value of each dimension to avoid the embedding being driven by the first few PCs.

We adopted a similar framework as Seurat v3^36^ for data integration by first identifying the mutual nearest neighbors (MNN) as anchors between datasets, and then aligning the datasets through the anchors.

To find anchors between snmC-seq and scRNA-seq, we first Z-score scaled the mCH matrix and expression matrix of CEGs across cells, and the resulting matrices are represented as *X* (mC cell-by-CEG) and *Y* (RNA cell-by-CEG), respectively. CCA was used to find the shared low dimensional embedding of the two datasets, solved by singular value decomposition (SVD) of their dot product *USV*^*T*^ *= XY*^*T*^. *U* and *V* were normalized by dividing the L2-norm of each row, and were used to find 5 MNNs as anchors and score anchors using the same method as Seurat v3.

The original CCA framework of Seurat (v3) is hard to scale up to millions of cells due to the memory bottleneck, where the mC cell-by-RNA matrix was used as the input to CCA. To handle this limitation, we randomly selected 50,000 cells from each dataset (*X*_*ref*_ and *Y*_*ref*_) as a reference to fit the CCA and transformed the other cells (*X*_*qry*_ and *Y*_*qry*_) onto the same CC space. Specifically, the canonical correlation vectors (CCV) of *X*_*ref*_ and *Y*_*ref*_ (denoted as *U*_*qry*_ and *V*_*qry*_) were computed by 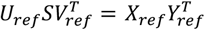 where 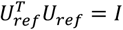 and 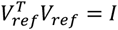. Then the CCV of *X*_*qry*_ and *Y*_*qry*_ (denoted as *U*_*qry*_ and *V*_*qry*_) were computed by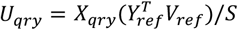 and 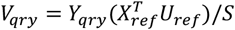. The embeddings from the reference and query cells were concatenated for anchor identification. To find anchors between snmC-seq and Epi-Retro-Seq, we used the snmC-seq data to fit a PCA model and use the model to transform Epi-Retro-Seq cells to the same space and find 5 nearest snmC-seq cells for each Epi-Retro-Seq cell. Reciprocally, we fit another PCA model with the Epi-Retro-Seq cells and transform the snmC-seq cells and find 5 nearest Epi-Retro-Seq cells for each snmC-seq cell. The mutual nearest neighbors between the two datasets were used as anchors and scored using the same method as Seurat v3.

The PCs derived from the previous step were then integrated together using the same method as Seurat v3 through these anchors. This integration step projects the PCs of Epi-Retro-Seq and scRNA-seq (query) to the PCs of the snmC-seq (reference) while keeping the PCs of the reference dataset unchanged. The resulting PCs from the three datasets were used for t-SNE visualization and k-nearest neighbor (k=25) graph construction with Euclidean distance. The joint clustering was performed with the Leiden algorithm on the graph using a resolution of 1.0.

The quality of the integration analysis was evaluated from two aspects. 1) We visualized the different modalities in the co-embedding space (**Extended Data Fig. 4 left**). The local neighborhoods of the coembedding usually contain cells from all modalities, suggesting a good mixture between the three datasets after integration. 2) We computed the proportion of cells in each mC cluster (**Extended Data Fig. 4 middle**) or RNA cluster (**Extended Data Fig. 4 right**) assigned to each cluster defined on the co-embedding space (co-cluster). Since we used the highest granularity of clustering from individual modalities (original cluster), the co-clusters were usually larger than the original clusters. We therefore used the proportion of original clusters rather than the proportion of co-clusters, to demonstrate that almost all original clusters are included in one co-cluster with low ambiguity. The strongest signals align on the diagonals suggesting that the co-embedding preserved the cluster structures that were originally present within each modality. Further evidence of integration quality was suggested by the downstream analyses, where highly consistent cell type specificity of marker gene expression and gene body mCH were observed (**Figs. 3f,g, 4e,f, and Extended Data Fig. 9a**).

### Cluster associated with projection

For neurons projecting to each target within one source, we computed the proportion of these neurons in each joint Leiden cluster. The clusters with >5% of the cells were considered as associated with the projection. The clusters associated with at least one projection were shown in the heatmaps of **Figs. 3, 4, 5**, and **Extended Data Fig. 5**. The values in the heatmaps represent the proportion of projection neurons in each cluster, Z-scored across the projection targets.

In general, there are two intuitive ways to quantify the enrichment of projection neurons in a cluster. One is to directly find the clusters with a high absolute proportion of Epi-Retro-Seq neurons projecting to a target. The other is to find clusters captured at a significantly higher frequency in the projection-enriched data relative to the unbiased data. The two methods each have their advantages and shortcomings. For example, the contaminated cells from inaccurate labeling or gating are likely to have similar distribution across clusters to unbiased profiling. So a comparison using unbiased data as a control might help exclude the contaminated clusters better. However, if most of the neurons from a projection type are in the clusters that are originally abundant cell types in the source, by comparing with unbiased data, we would miss the predominant clusters making the projection. In this manuscript, we used the absolute proportions but not the relative ones to the unbiased data due to the different profiling strategies between the two datasets. Although the Epi-Retro-Seq samples and unbiased snmC-seq samples were dissected in the same way, we pooled the different dissections into the 32 different sources to perform FANS and sequencing for Epi-Retro-Seq, so that the proportion of cells from different dissection regions of the same source is likely to follow their proportions in the mouse brain. However, the unbiased snmC-seq profiled all the dissection regions separately and sequenced the same number of cells in each dissection, which manually amplified the proportion of cells from the smaller/sparser dissection regions relative to the larger/denser ones, and limited the power to estimate the real proportion of neurons in each cluster from the sources.

### Classification of MERFISH cells into major brain regions and cell clusters

The MERFISH experiments were conducted as described by Liu et al. (companion paper #6), including the gene panel design, tissue preparation, imaging, data processing, and annotation. The dataset includes two sagittal slices (S1 and S2, where S1 is more lateral and S2 is more medial) and 14 coronal slices (C2, C4, C6, C8, C10, C12, and C14, roughly corresponding to slice 2, 4, 6, 8, 10, 12, and 14 in **Extended Data Fig. 1**, with two replicates for each slice, represented as R1 and R2). The same naming of slices was used throughout this manuscript (**Figs. 3e, 4d, 5c and Extended Data Fig. 6**).

The MERFISH cells were classified into subclasses and brain region groups by integration with scRNA-seq data. The 489 autosomal genes overlapped between scRNA-seq and MERFISH datasets were used. We fit a PCA model with the scRNA-seq cells and transformed the MERFISH cells with the model. The PCs were normalized by the singular value of each dimension. The cell-by-gene matrices were Z-score normalized across cells within each dataset, and CCA was used to find anchors between the two datasets. We used 50,000 cells to fit the CCA and transformed the other cells as described above. The transformed PCs of MERFISH cells were then aligned to the PCs of scRNA-seq cells to derive a coembedding between the two datasets. This co-embedding was used for label transfer of cell subclasses from scRNA-seq data to MERFISH data, considering 25 neighboring scRNA-seq cells for each MERFISH cell.

The cells classified as non-neuronal and immature neuronal subclasses were excluded due to lack of regional specificity, and the rest of cells from the two datasets were integrated again with the procedures described above to transfer the label of 14 brain region groups from the scRNA-seq neurons to the MERFISH neurons. The initial label assignment is noisy. Therefore a smoothing step was performed to refine the region group assignment. Specifically, for each MERFISH cell i, we found its 25 neighbors on the same slice (denoted as *Ns*_*i*_) based on the spatial coordinates, and used *D s*_*i*_ to represent the corresponding distances between i and its j-th neighbor *Ns*_*i,j*_. Similarly, we found the 25 scRNA-seq neighbors for each MERFISH cell *i* based on the integration, and used *D r Ns*_*i*_ to represent the distances. The distance matrices were transformed as described in the label transfer section, and the final spatial labels were transferred from the 25 RNA neighbors of each of the 25 spatial neighbors (625 scRNA-seq cells in total) to one MERFISH cell. The weight between the MERFISH cell i and the k-th scRNA-seq neighbor of its j-th spatial neighbor was computed as 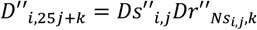 is row normalized and used as weights for label transfer as described in previous sections.

We note that this could also be achieved through registration of MERFISH DAPI images to the common coordinate framework. However, our companion works demonstrated that the procedure is relatively challenging and it is important to use cell types with known locations as landmarks to refine the registration.

Finally, the neurons from each region group were selected and integrated with the scRNA-seq cells from the same region group, using the same procedure as described above. The methylation-RNA co-cluster labels were transferred from the scRNA-seq cell to the MERFISH cells. The MERFISH cells assigned to the MM and DCO subclasses in the last step were also excluded since those clusters were not included in the co-cluster analysis as described in the previous section.

### Comparison with Act-Seq

The 10x scRNA-seq data in Kim et al.^6^ were downloaded from Mendeley Data (https://data.mendeley.com/datasets/ypx3sw2f7c/3), and are referred to as Act-Seq in this section and **Extended Data Fig. 7**. The dataset contains 168,877 cells in total, among which 78,476 were labeled as neurons and were used to integrate to the unbiased scRNA-seq data of hypothalamic neurons, using the 5,314 CEGs of 1,891 L3 scRNA-seq clusters. PCA was fitted with scRNA-seq data and the Act-Seq data was transformed. The CCA framework was used to find anchors between the two datasets, and the transformed PCs of Act-Seq were aligned to the scRNA-seq PCs. We used the label transfer method described in the previous section to transfer the mC-RNA co-cluster labels from scRNA-seq cells to the Act-Seq cells, considering 5 neighboring scRNA-seq cells for each Act-Seq cell. The Act-Seq cells with *Fos* expression level >0 were considered to be *Fos*+ cells, and the proportion of *Fos*+ cells were compared between control and each behavior using Fisher exact tests. The Fos expression levels were compared with Wilcoxon rank-sum tests. Only the 23,345 Act-Seq cells from 16 VMH neuron clusters were considered as VMHvl cells and were used for the behavior association studies in Kim et al. However, almost none of them correspond to projection-associated clusters in our data (**Extended Data Fig. 7a-c**). We further compared our projection-associated clusters with all the neuron clusters profiled in Kim et al. and note that five clusters have corresponding clusters in the Act-Seq data (**Extended Data Fig. 7d, in red**). Among them, clusters 4 and 64 showed weak but significant increases in proportions of *Fos*+ cells labeled during certain behaviors (**Extended Data Fig. 7f**).

The generally weak associations between projection-associated and behavior-associated clusters are likely due to the small overlap between the brain regions profiled in the two datasets, particularly the under representation of VMHvl neurons in Epi-Retro-Seq data. Additionally, because there were far fewer cells profiled in Epi-Retro-Seq vs. Act-Seq, the granularity of clusters used for projection-association and behavior association is different; this difference is particularly pronounced in VMHvl where >10 times more cells were used for the behavior association study (**Extended Data Fig. 7c-e**). Therefore, further increasing the size of datasets to achieve higher granularity of cell typing in specific regions of interest could facilitate further association between molecular types with projections and behaviors. Our study aimed at a comprehensive view of a large number of projections across the whole brain and focused on targets that do not appear to receive strong input from VMH. This apparently limited the data overlap between these and limited the ability to make direct comparisons between studies.

### Comparison with Retro-Seq

The scRNA-seq data in Phillips et al.^7^ were downloaded from the Gene Expression Omnibus with the identifier GSE133912. The Retro-Seq data were integrated to the unbiased scRNA-seq data of thalamic neurons, using the 5,404 CEGs of 1,128 L3 scRNA-seq clusters. PCA was fitted with scRNA-seq data and Retro-Seq data was transformed. The CCA framework was used to find anchors between the two datasets, and the transformed PCs of Retro-Seq were aligned to the scRNA-seq PCs. To compare the distribution of Retro-Seq cells and Epi-Retro-Seq cells across thalamic cell clusters, we used the label transfer method described in the previous section to transfer the mC-RNA co-cluster label and the joint t-SNE coordinates from scRNA-seq cells to the Retro-Seq cells, considering 5 neighboring scRNA-seq cells for each Retro-Seq cell.

### Differentially expressed genes

The gene expression level of each single cell was normalized by the total UMI count of the cell and log-transformed. We performed pairwise comparisons between clusters associated with projection neurons. For each cluster pair, the *P* values were derived with the Wilcoxon rank-sum test, and the fold-change is computed as the ratio between the average expression level across cells in the two clusters. The genes with absolute value of log2 fold-change greater than 1 and False Discovery Rate (FDR, Benjamini-Hochberg Procedure) values smaller than 0.01 were considered as differentially expressed. The DEGs from all cluster pairs were merged to generate the heatmaps in **Fig. 3** and **Fig. 4**. Only the top 100 DEGs ranked by FDRs were used if there were more than 100 DEGs identified between a pair of clusters.

### Differentially methylated regions (DMRs) and association with genes

The unbiased snmC-seq cells from each mC-RNA co-cluster were merged to generate pseudobulk methylation profiles. The Epi-Retro-Seq cells were not used due to the different genome backgrounds of the mice to avoid confounding results. DMRs were identified within each region group between clusters using ALLCools. We then calculated the Pearson Correlation Coefficient (PCC) between DMR mCG and gene mCH fraction. For a group of overlapping DMRs, we selected the one with the highest absolute PCC value to represent that group, making the edges’ DMRs non-overlap. Similar to the domain boundary and interaction correlation analysis, we shuffled the DMRs and genes within each sample to calculate null PCC and estimate FDR. We filtered DMR-Target edges with FDR < 0.001.

### Transcription factor motif enrichment

We used an ensemble motif database from SCENIC+^39^, which contained 49,504 motif position weight matrices (PWM) collected from 29 sources. Redundant motifs (highly similar PWMs) were combined into 8,045 motif clusters through clustering based on PWM distances calculated by TOMTOM^40^ by the SCENIC+ authors. Each motif cluster was annotated with one or more mouse TF genes. To calculate motif occurrence on DMRs, we used the Cluster-Buster^41^ implementation in SCENIC+, which scanned motifs in the same cluster together with Hidden Markov Models. Within each region group, we assign hypo-DMRs to each cluster if the mCG level of a DMR in the cluster is below the 10th quantile of all DMRs from the region group and below the 10th quantile of the mCG level of this DMR in all clusters from the region group. To perform motif enrichment analysis, we used the recovery-curve-based cisTarget algorithm^39^. In brief, the cisTarget algorithm performed motif enrichment on the hypo-DMRs of each cluster by calculating the area under the recovery curve (AUC) for each motif, which is further normalized based on all other motifs in the collection to calculate a Normalized Enrichment Score (NES). We used the cutoff AUC>0.01 and NES > 3 to select enriched motifs. The TFs shown in **Fig. 3i** and **Fig. 4h** were additionally required to have expression level >0 and normalized mCH level <1 in at least one cluster that its motif enriched in, to select the TFs that are likely to express among a family of TFs showing the same motif enrichment scores.

## Data access and code availability

The data can be accessed via the NeMO ftp archive: https://data.nemoarchive.org/biccn/grant/u19_cemba/ecker/epigenome/sncell/mCseq2_retro/mouse/. Raw and processed data are also available at GEO under accession code GSE230782. The code for all of the analyses can be found at https://github.com/zhoujt1994/EpiRetroSeq2023.git.

## Author contribution

Contribution to research design: E.M.C., Z.Z, M.M.B., J.R.E., J.Z., X.J., K.L.

Contribution to data collection: Z.Z., Y.P., A.B., A.R., W.N.L., E.W., C.L., P.A.M, A.A., J.L.A.S., N.C., M.L., L.B., C.F., Z.Y., K.A.S., B.T.,

J.A., M.A.K., C.V., J.R.N., R.G.C., N.S.P., M.V., M.R., M.J., T.I., J.O.,

N.E., J.L., S.C., J.R., S.H., A.P.-D., B.D., J.B.S, C.O., H.Z., M.M.B.

Contribution to data analysis: J.Z., Z.Z., E.M.C, M.W., H.L., Q.Z., Z.Y., H.Z.

Contribution to data archive/infrastructure: E.A.M., J.Z., Z.Z., Y.P., A.B., A.R.

Contribution to research coordination: Z.Z., E.M.C., M.M.B., J.R.E., J.Z., Y.P., X.J., E.W., C.L., E.A.M., K.L.

Contribution to writing manuscript: J.Z., Z.Z., E.M.C., J.R.E.

## Acknowledgements

We are grateful to Dr. Michael Nunn for help with management of the project. This work is supported by NIMH U19MH114831 to E.M.C and J.R.E. under the BRAIN Initiative of National Institutes of Health (NIH) and by NEI R01EY022577 to E.M.C. The Flow Cytometry Core Facility of the Salk Institute (RRID:SCR_014839) is supported by funding from NIH-NCI CCSG: P30 014195 and Shared Instrumentation Grant S10-OD023689 (Aria Fusion cell sorter). J.R.E is an investigator of the Howard Hughes Medical Institute. The 10x sn/scRNA-seq was funded by the U19MH114830 grant to H.Z. and X.Z., under the BRAIN Initiative of National Institutes of Health (NIH).

## Extended data figures

**Extended Data Figure 1.**
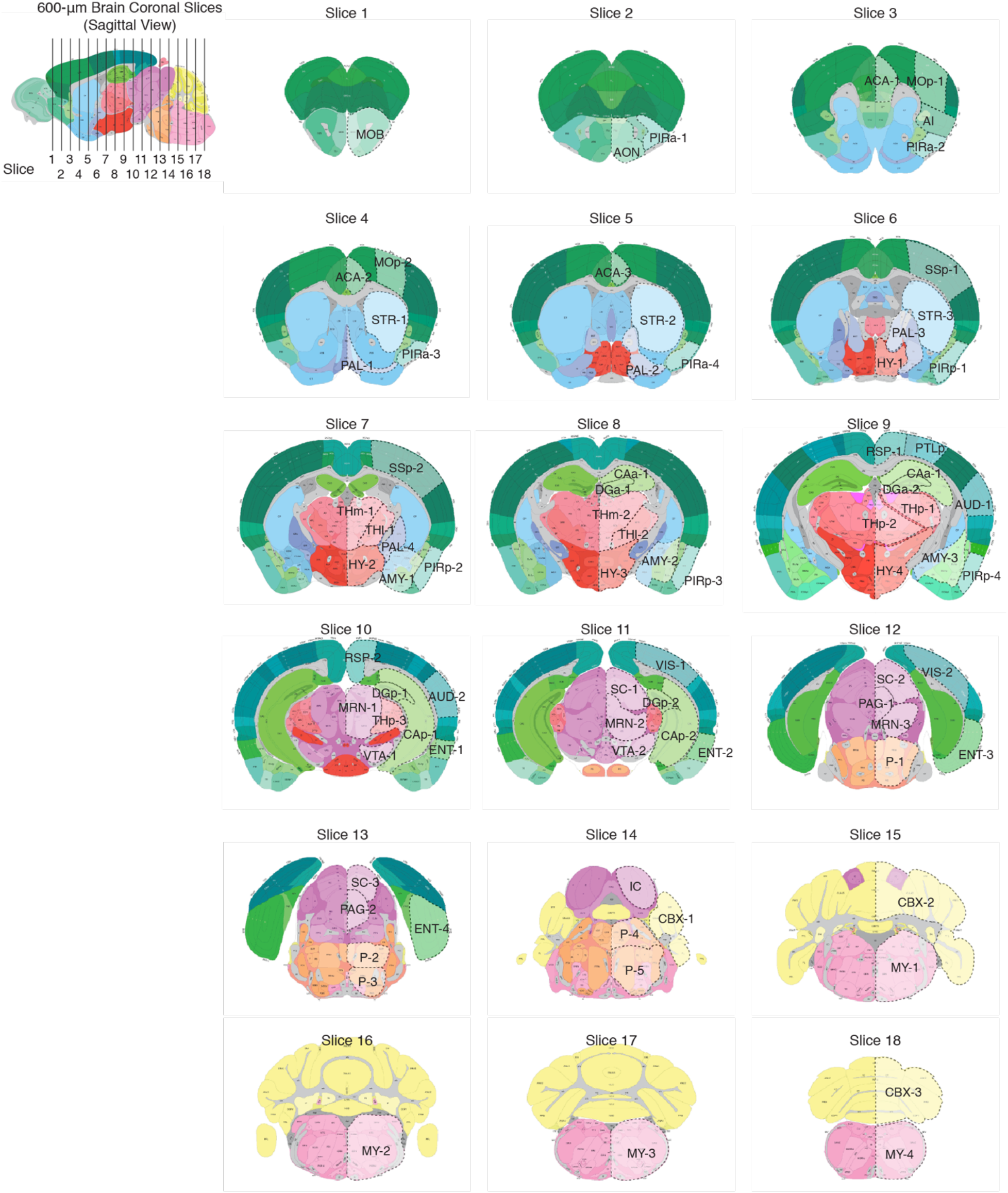
The dissection map of source regions across the mouse brain. The posterior views of dissected slices are shown. The slices correspond to Allen Mouse Common Coordinate Framework (CCF), Reference Atlas, Version 3 (2020), level 21∼27 (slice 1), 27∼33 (slice 2), 33∼39 (slice 3), 39∼45 (slice 4), 45∼51 (slice 5), 51∼57 (slice 6), 57∼63 (slice 7), 63∼69 (slice 8), 69∼75 (slice 9), 75∼81 (slice 10), 81∼87 (slice 11), 87∼93 (slice 12), 93∼99 (slice 13), 99∼105 (slice 14), 105∼111 (slice 15), 111∼117 (slice 16), 117∼123 (slice 17), and 123∼129 (slice 18), respectively. Regions dissected from each slice are indicated by dotted lines and are annotated.

**Extended Data Figure 2.**
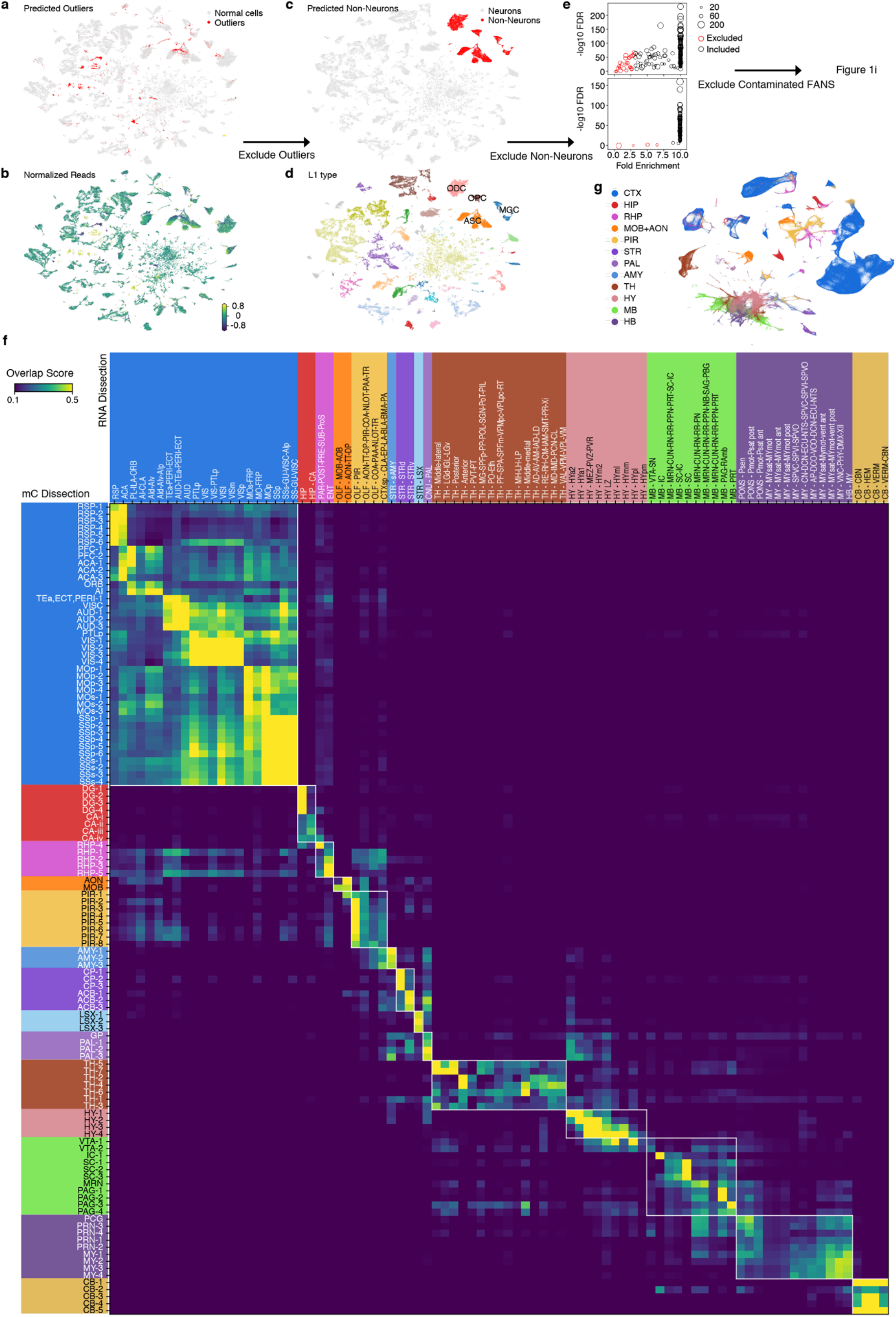
Quality control workflow and region group assignment. **a, b**, Joint t-SNE of Epi-Retro-Seq cells (n=56,843) and unbiased snmC-seq cells (n=310,605) after basic QC (Methods, QC Step 1) colored by the predicted outliers (**a**) or the total number of reads normalized per sequencing plate (**b**). **c, d**, Joint t-SNE of Epi-Retro-Seq cells (n=48,032) and unbiased snmC-seq cells (n=301,626) after removing outlier clusters (Methods, QC Step 2) colored by neuronal vs. non-neuronal cells (**c**) or their assigned L1 type (**d**). **e**, the on-target vs. off-target fold enrichment (x-axis) and -log10 FDR (y-axis) of IT (n=186, top) or ET (n=100, bottom) FANS experiments. The size of the circle is proportional to the number of neurons captured in the experiment. **f**, The overlap scores between 115 snmC-seq dissections and 87 scRNA-seq dissections. Each region group is colored differently on the x and y axes and squared in the heatmap. **g**, Joint t-SNE of Epi-Retro-Seq (n=35,743), snmC-seq (n=266,740), and scRNA-seq (n=2,434,472) neurons colored by region groups. (**f)** and (**g)** share the same color palette.

**Extended Data Figure 3.**
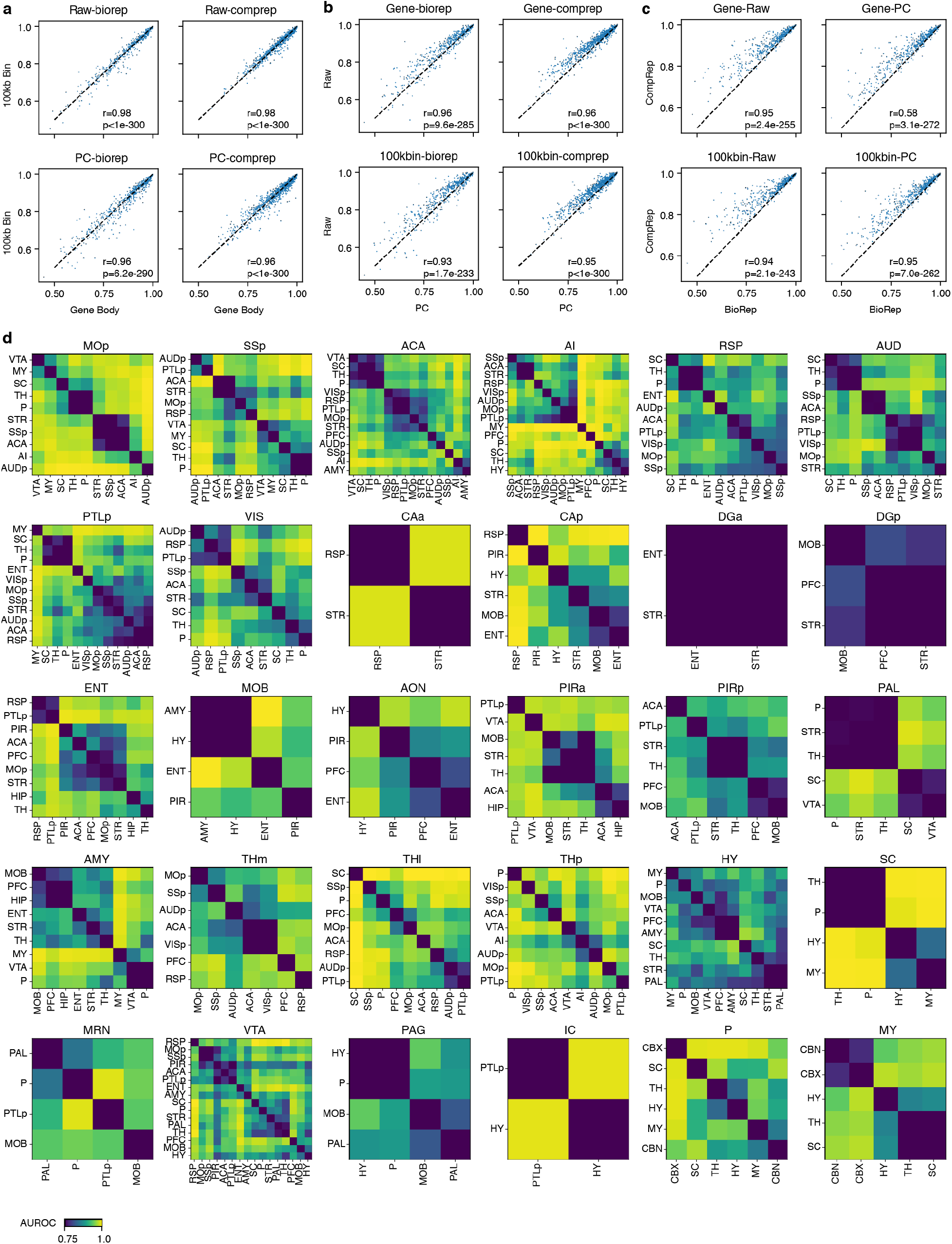
Quantification of target discriminability for all 926 target pairs from all source regions. **a**, Comparisons of AUROCs from models trained with gene features (x-axis) or 100kb bin features (y-axis) using posterior mCH level (top) or mCH principal components (bottom) when splitting the training and testing data according to biological replicates (left) or computational replicates (right). **b**, Comparisons of AUROCs from models trained with mCH principal components (x-axis) or posterior mCH level (y-axis) using gene (top) or 100kb bin (bottom) features when split the training and testing data according to biological replicates (left) or computational replicates (right). **c**, Comparisons of AUROCs from models when splitting the training and testing data according to biological replicates (x-axis) or computational replicates (y-axis) using gene (top) or 100kb bin (bottom) features and posterior mCH level (left) or mCH principal components (right). For **a-c**, each dot represents a pairwise comparison between the neurons projecting from the same source to two different targets. The plots involving biological replicates have 516 data points each while the others have 926 data points each. Pearson Correlation Coefficient (PCC, r) and *P* value (permutation test) are labeled in each panel. **d**, The AUROC between neurons projecting from each of the 30 sources to all possible pairs of targets that have been profiled for the source. STR and CBX are not included since we only profiled one target for these sources.

**Extended Data Figure 4.**
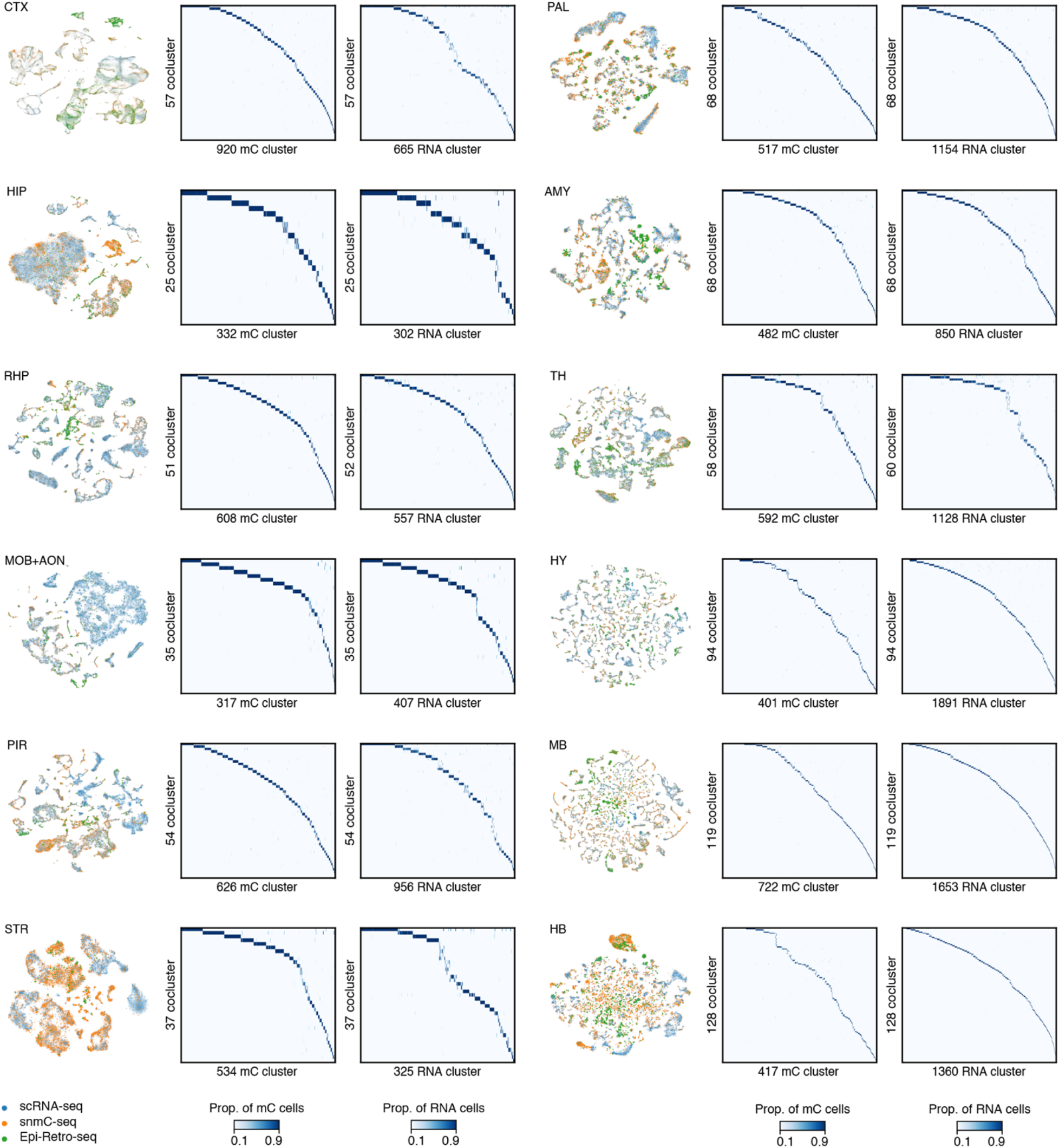
Co-clustering of Epi-Retro-Seq, unbiased snmC-seq and scRNA-seq data for 12 brain regions. For each brain region group, the joint t-SNE colored by data modality (left) and the proportion of cells from each snmC-seq L4 cluster (middle, column) or scRNA-seq L3 cluster (right, column) within each co-cluster (middle and right, row). Numbers of co-clusters in the rows are sometimes different for middle and right columns because only the co-clusters (rows) with more than one value >10% across the columns are shown. See **Methods** for further details.

**Extended Data Figure 5.**
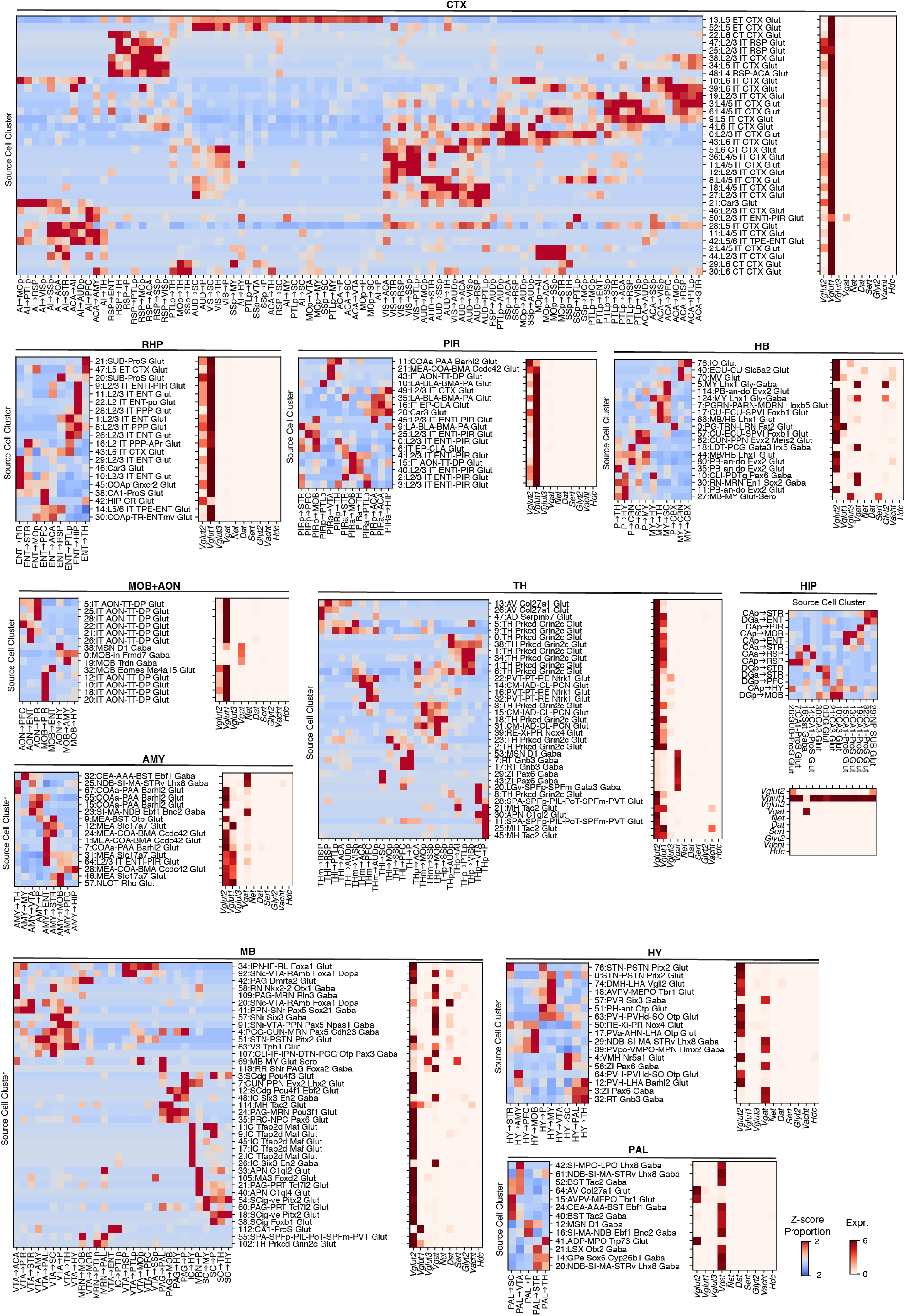
Projection-enriched cell clusters and their neurotransmitter usage for all brain regions. Joint clustering analysis of Epi-Retro-Seq, unbiased snmC-seq and scRNA-seq was performed on each of the major brain region groups, including CTX, RHP, PIR, HB, MOB+AON, AMY, TH, HIP, MB, HY, and PAL (STR not included because there is only one target), to characterize neuronal cell clusters that were enriched for Epi-Retro-Seq projections. The normalized proportion of each projection in each cluster was visualized in the heatmaps (left) for each of the brain region groups. In addition, the expression levels of 10 marker genes for neurotransmitter usage in each cluster are visualized in the heatmap (right) for each brain region group. These genes included *Slc17a7* (*Vglut1*), *Slc17a6* (*Vglut2*), and *Slc17a8* (*Vglut3*) for glutamatergic neurons, *Slc32a1* (*Vgat*) for GABAergic neurons, *Slc6a2* (*Net*) for noradrenergic neurons, *Slc6a3* (*Dat)* for dopaminergic neurons, *Slc6a4* (*Sert*) for serotonergic neurons, *Slc6a5* (*Glyt2*) for glycinergic neurons, *Slc18a3* (*Vacht*) for cholinergic neurons, and histidine decarboxylase (*Hdc*) for histaminergic neurons.

**Extended Data Figure 6.**
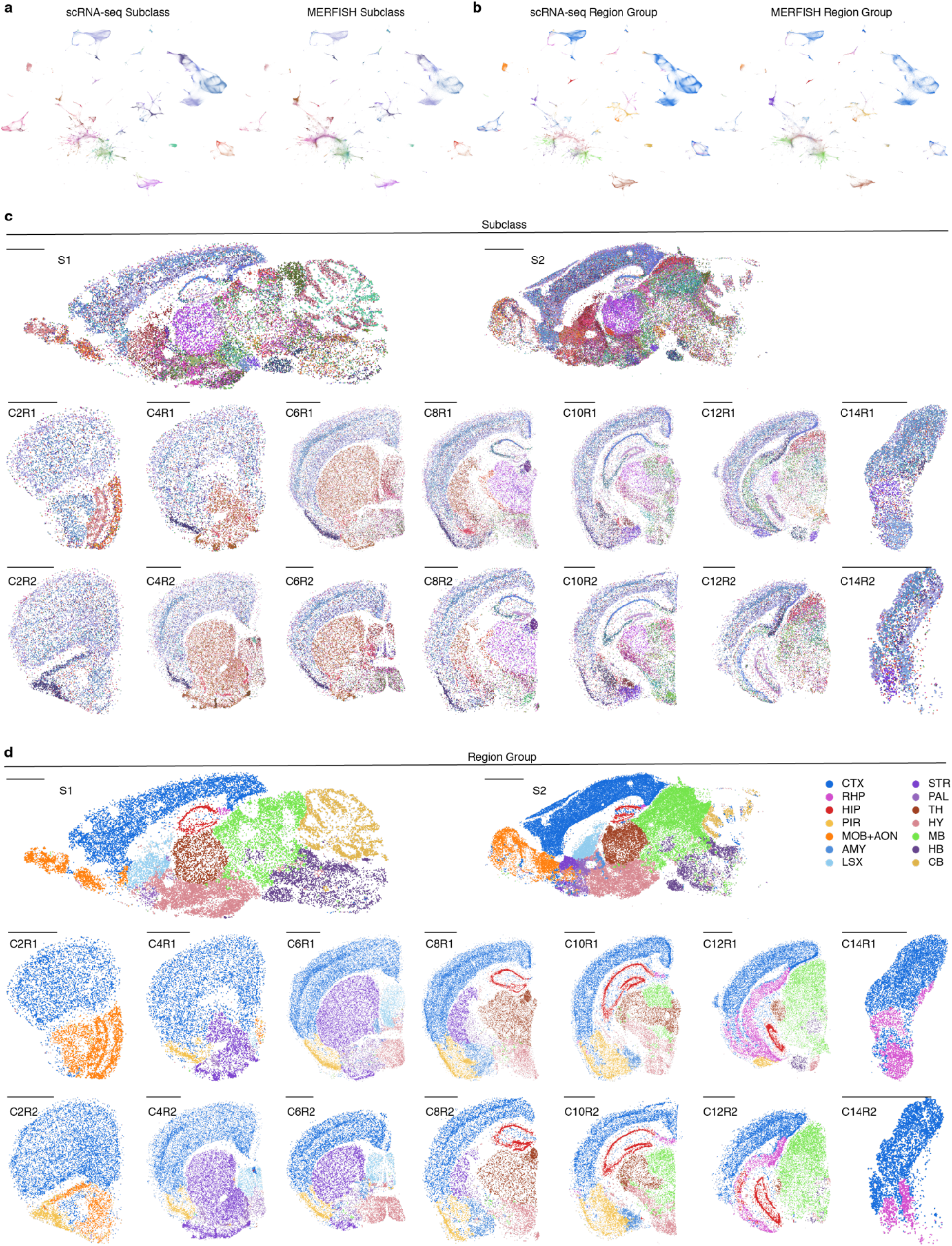
Joint clustering and annotation of MERFISH and scRNA-seq data. **a, b**, Joint t-SNE of scRNA-seq neurons (n=2,619,158, left) and MERFISH neurons (n=329,282, right) colored by cell subclass (**a**) or region group (**b**). The labels for scRNA-seq cells are based on Yao et al.^16^ and the labels for MERFISH cells are predicted through integration. **c, d**, The MERFISH slices colored by cell subclass (**c**) or region group (**d**). Scale bars represent 15 mm.

**Extended Data Figure 7.**
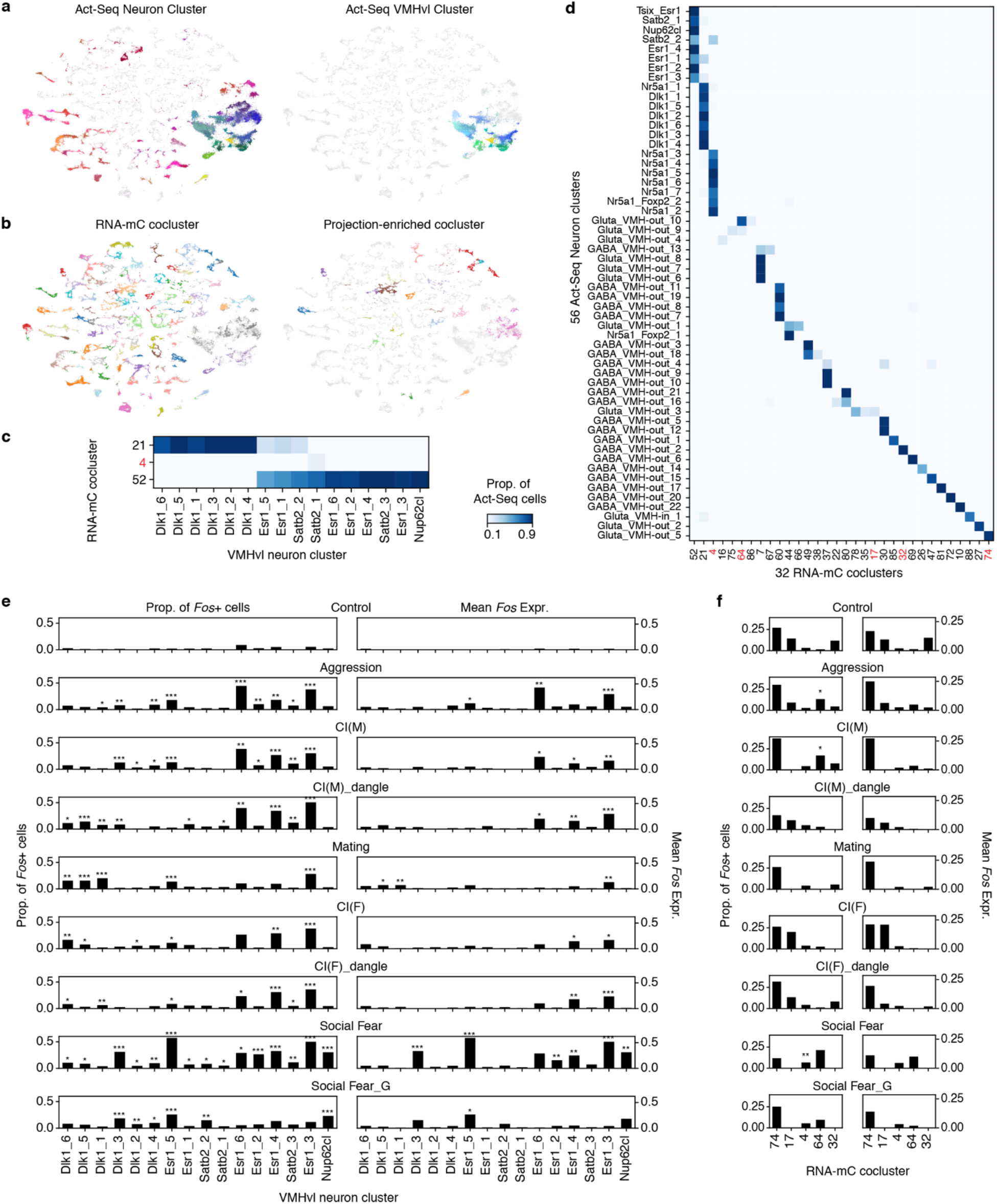
Integration and comparisons of hypothalamic Epi-Retro-Seq, Act-seq, and scRNAseq data. **a, b**, Joint t-SNE of Act-Seq neurons^6^ (n=78,476, **a**) and scRNA-seq neurons (n=148,840, **b**). In (**a**), all Act-Seq neurons (left) or only VMHvl neurons (right) are colored by Act-Seq neuron cluster (left) or VMH cluster (right). In (**b**), all scRNA-seq neurons (left) or only neurons in projection associated clusters (right) are colored by the co-cluster label. **c, d**, The proportion of neurons from each of the Act-Seq VMHvl clusters (**c**) or all neuron clusters (**d**) classified as neurons of each co-cluster. Only the co-clusters with value >0.1 in at least one Act-Seq cluster are shown. The projection-associated co-clusters are labeled in red. **e, f**, Proportion of Fos+ “behavior activated” cells (left) or average Fos expression (right) of each VMHvl cluster (**e**) or each co-cluster (**f**) in control and different behavior experiments. Only the co-clusters labeled red in d are shown in (**f**). ^*, **^, and ^***^ represent FDR<0.1, 0.01, and 0.001, respectively.

**Extended Data Figure 8.**
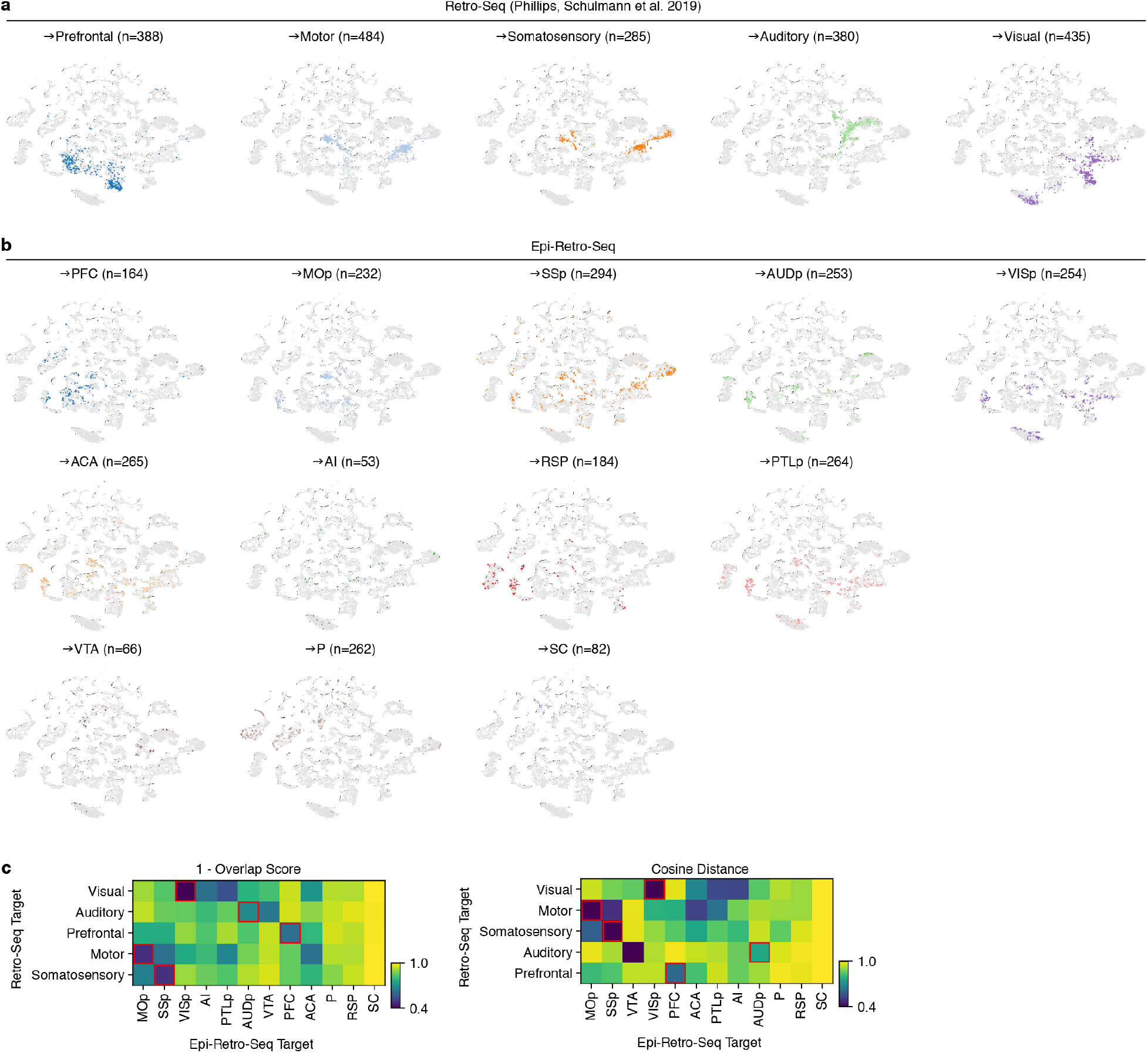
Comparison of thalamic Retro-Seq and Epi-Retro-Seq data. **a**, t-SNEs for visualization of the Retro-Seq data from thalamic neurons projecting to prefrontal, motor, somatosensory, auditory, and visual cortices^7^ that were mapped onto the joint-clustering analysis of Epi-Retro-Seq, unbiased snmC-seq and scRNA-seq in TH. **b**, The t-SNEs for visualization of the Epi-Retro-Seq data for thalamic neurons projecting to 12 different targets that were mapped onto the same t-SNE space. **c**, The overlap score and cosine distance were calculated for each pairwise comparison of Retro-Seq and Epi-Retro-Seq projections and were visualized in the heatmaps, respectively.

**Extended Data Figure 9.**
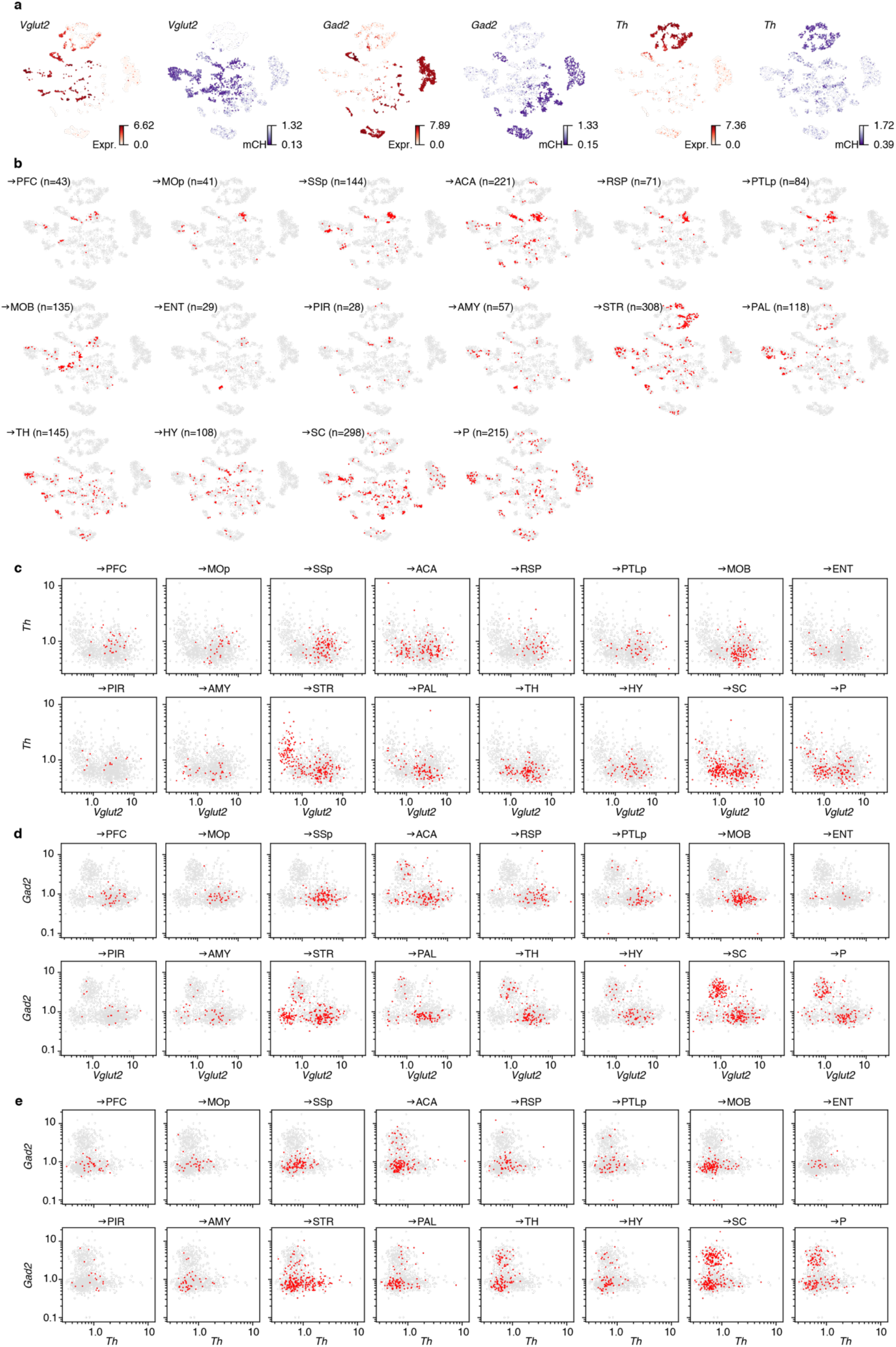
The neurotransmitter usage of VTA projection neurons. **a**, Joint t-SNE of Epi-Retro-Seq, unbiased snmC-seq and scRNA-seq of VTA neurons colored by the gene expression levels (red) and gene-body mCH levels (purple) for *Vglut2* (left), *Gad2* (middle), and *Th* (right), marker genes for glutamatergic, GABAergic, and dopaminergic neurons, respectively. **b**, The distribution of VTA neurons projecting to each of the 16 targets on the same t-SNE. **c-e**, The gene-body mCH levels of *Th* versus *Vglut2* (**c**), *Gad2* versus *Vglut2* (**d**), and *Gad2* versus *Th* (**e**) for VTA neurons projecting to each of the 16 targets, are visualized in scatter plots. Note that, because low mCH levels indicate high gene expression, the axes in **c-e** are plotted as the reciprocal mCH values (1/gene body mCH), so low mCH is plotted to the right/up and high to the left/down.

## Supplementary Tables

**Supplementary Table 1. Epi-Retro-Seq injection information**.

**Supplementary Table 2. Epi-Retro-Seq cell metadata**.

**Supplementary Table 3. Gene categories for ROC analyses**.

**Supplementary Table 4. Comparisons of injection coordinates to retrogradely label thalamic projection neurons for Epi-Retro-Seq vs. Retro-Seq**.

## References

1. Chen, X. et al. High-Throughput Mapping of Long-Range Neuronal Projection Using In Situ Sequencing. Cell 179, 772–786.e19 (2019).

2. Sun, Y.-C. et al. Integrating barcoded neuroanatomy with spatial transcriptional profiling enables identification of gene correlates of projections. Nat. Neurosci. 24, 873–885 (2021).

3. Economo, M. N. et al. Distinct descending motor cortex pathways and their roles in movement. Nature 563, 79–84 (2018).

4. Kim, E. J. et al. Extraction of Distinct Neuronal Cell Types from within a Genetically Continuous Population. Neuron 107, 274–282.e6 (2020).

5. Zhang, Z. et al. Epigenomic diversity of cortical projection neurons in the mouse brain. Nature 598, 167–173 (2021).

6. Kim, D.-W. et al. Multimodal Analysis of Cell Types in a Hypothalamic Node Controlling Social Behavior. Cell 179, 713–728.e17 (2019).

7. Phillips, J. W. et al. A repeated molecular architecture across thalamic pathways. Nat. Neurosci. 22, 1925–1935 (2019).

8. Mo, A. et al. Epigenomic Signatures of Neuronal Diversity in the Mammalian Brain. Neuron 86, 1369–1384 (2015).

9. Luo, C. et al. Single nucleus multi-omics identifies human cortical cell regulatory genome diversity. Cell Genom 2, p(2022).

10. Lister, R. et al. Global epigenomic reconfiguration during mammalian brain development. Science 341, 1237905 (2013).

11. Tervo, D. G. R. et al. A Designer AAV Variant Permits Efficient Retrograde Access to Projection Neurons. Neuron 92, 372–382 (2016).

12. Wang, Q. et al. The Allen Mouse Brain Common Coordinate Framework: A 3D Reference Atlas. Cell 181, 936–953.e20 (2020).

13. Luo, C. et al. Single-cell methylomes identify neuronal subtypes and regulatory elements in mammalian cortex. Science 357, 600–604 (2017).

14. Luo, C. et al. Robust single-cell DNA methylome profiling with snmC-seq2. Nat. Commun. 9, 3824 (2018).

15. Tian, W. et al. Epigenomic complexity of the human brain revealed by single-cell DNA methylomes and 3D genome structures. bioRxiv 2022.11.30.518285 (2022) doi:10.1101/2022.11.30.518285.

16. Yao, Z. et al. A high-resolution transcriptomic and spatial atlas of cell types in the whole mouse brain. bioRxiv 2023.03.06.531121 (2023) doi:10.1101/2023.03.06.531121.

17. Zhang, M. et al. A molecularly defined and spatially resolved cell atlas of the whole mouse brain. bioRxiv 2023.03.06.531348 (2023) doi:10.1101/2023.03.06.531348.

18. Benevento, M., Hökfelt, T. & Harkany, T. Ontogenetic rules for the molecular diversification of hypothalamic neurons. Nat. Rev. Neurosci. 23, 611–627 (2022).

19. Moffitt, J. R. et al. Molecular, spatial, and functional single-cell profiling of the hypothalamic preoptic region. Science 362, eaau5324 (2018).

20. Martin Usrey, W. & Murray Sherman, S. Cell Types in the Thalamus and Cortex. in (Oxford University Press, 2021).

21. Sripanidkulchai, K. & Wyss, J. M. Thalamic projections to retrosplenial cortex in the rat. J. Comp. Neurol. 254, 143–165 (1986).

22. Zander, J.-F. et al. Synaptic and vesicular coexistence of VGLUT and VGAT in selected excitatory and inhibitory synapses. J. Neurosci. 30, 7634–7645 (2010).

23. Kramer, S. G., Kidd, T., Simpson, J. H. & Goodman, C. S. Switching repulsion to attraction: changing responses to slit during transition in mesoderm migration. Science 292, 737–740 (2001).

24. Hohenester, E., Hussain, S. & Howitt, J. A. Interaction of the guidance molecule Slit with cellular receptors. Biochem. Soc. Trans. 34, 418–421 (2006).

25. Farghaian, H. et al. Scapinin-induced inhibition of axon elongation is attenuated by phosphorylation and translocation to the cytoplasm. J. Biol. Chem. 286, 19724–19734 (2011).

26. Miyata, T. et al. Neuron-enriched phosphatase and actin regulator 3 (Phactr3)/ nuclear scaffold-associated PP1-inhibiting protein (Scapinin) regulates dendritic morphology via its protein phosphatase 1-binding domain. Biochem. Biophys. Res. Commun. 528, 322–329 (2020).

27. Trudeau, L.-E. et al. The multilingual nature of dopamine neurons. Prog. Brain Res. 211, 141–164 (2014).

28. Bouarab, C., Thompson, B. & Polter, A. M. VTA GABA Neurons at the Interface of Stress and Reward. Front. Neural Circuits 13, 78 (2019).

29. Cai, J. & Tong, Q. Anatomy and Function of Ventral Tegmental Area Glutamate Neurons. Front. Neural Circuits 16, 867053 (2022).

30. Phillips, R. A., 3rd et al. An atlas of transcriptionally defined cell populations in the rat ventral tegmental area. Cell Rep. 39, 110616 (2022).

31. Yamaguchi, T., Sheen, W. & Morales, M. Glutamatergic neurons are present in the rat ventral tegmental area. Eur. J. Neurosci. 25, 106–118 (2007).

32. Yamaguchi, T., Wang, H.-L., Li, X., Ng, T. H. & Morales, M. Mesocorticolimbic glutamatergic pathway. J. Neurosci. 31, 8476–8490 (2011).

33. Lacar, B. et al. Nuclear RNA-seq of single neurons reveals molecular signatures of activation. Nat. Commun. 7, 11022 (2016).

34. Liu, H. et al. DNA methylation atlas of the mouse brain at single-cell resolution. Nature 598, 120– 128 (2021).

35. Miles, A. et al. zarr-developers/zarr-python: v2.5.0. (2020). doi:10.5281/zenodo.4069231.

36. Stuart, T. et al. Comprehensive Integration of Single-Cell Data. Cell 177, 1888–1902.e21 (2019).

37. Smith, S. J. et al. Single-cell transcriptomic evidence for dense intracortical neuropeptide networks. Elife 8, p(2019).

38. Tasic, B. et al. Adult mouse cortical cell taxonomy revealed by single cell transcriptomics. Nat. Neurosci. 19, 335–346 (2016).

39. González-Blas, C. B. et al. SCENIC+: single-cell multiomic inference of enhancers and gene regulatory networks. bioRxiv 2022.08.19.504505 (2022) doi:10.1101/2022.08.19.504505.

40. Gupta, S., Stamatoyannopoulos, J. A., Bailey, T. L. & Noble, W. S. Quantifying similarity between motifs. Genome Biol. 8, R24 (2007).

41. Frith, M. C., Li, M. C. & Weng, Z. Cluster-Buster: Finding dense clusters of motifs in DNA sequences. Nucleic Acids Res. 31, 3666–3668 (2003).

